# Genetic hypogonadal (Gnrh1^hpg^) mouse model uncovers influence of reproductive axis on maturation of the gut microbiome during puberty

**DOI:** 10.1101/2024.07.01.601610

**Authors:** Laura Sisk-Hackworth, Shayan R. Akhavan, Dennis D. Krutkin, Scott T. Kelley, Varykina G. Thackray

## Abstract

The gut microbiome plays a key role in human health and gut dysbiosis is linked to many sex-specific diseases including autoimmune, metabolic, and neurological disorders. Activation of the hypothalamic-pituitary-gonadal (HPG) axis during puberty leads to sexual maturation and development of sex differences through the action of gonadal sex steroids. While the gut microbiome also undergoes sex differentiation, the mechanisms involved remain poorly understood. Using a genetic hypogonadal (hpg) mouse model, we sampled the fecal microbiome of male and female wild-type and hpg mutant mice before and after puberty to determine how microbial taxonomy and function are influenced by age, sex, and the HPG axis. We showed that HPG axis activation during puberty is required for sexual maturation of the gut microbiota composition, community structure, and metabolic functions. We also demonstrated that some sex differences in taxonomic composition and amine metabolism developed independently of the HPG axis, indicating that sex chromosomes are sufficient for certain sex differences in the gut microbiome. In addition, we showed that age, independent of HPG axis activation, led to some aspects of pubertal maturation of the gut microbiota community composition and putative functions. These results have implications for microbiome-based treatments, indicating that sex, hormonal status, and age should be considered when designing microbiome-based therapeutics.

## INTRODUCTION

Studying sex differences is important for understanding and developing treatments for diseases with a sex bias. Many diseases are sex-biased in their incidence, prevalence, and/or severity, including numerous autoimmune[1], cardiovascular [2], and metabolic diseases [3]. Additionally, women are more likely to experience adverse drug reactions, due to both the historical exclusion of women from clinical trials and sex differences in drug pharmacokinetics [4–6].

Sex differences in mammals arise from sex chromosome genes and gonadal sex steroids. Differences in genetic content and gene doses between XX and XY individuals lead to sex differences in gene expression [7]. Though most X genes are inactivated, about 15% of genes escape inactivation, resulting in double gene dosage for those genes in females compared to males [8]. X chromosome gene dosage has been shown to impact immunity, inflammation, and adiposity and is associated with rates of stroke severity and the autoimmune disorder, systemic lupus erythematosus; two disorders with sex bias [1, 9–11]. The Y chromosome contains fewer genes than the X, but still contributes to development of sex differences beyond the SRY gene that leads to formation of the testes during gestation. Y chromosome genes are primarily expressed in the testes or are housekeeping genes homologous to X chromosome genes [12]. Sex differences in expression of X/Y chromosome homologous genes have been observed in nonreproductive tissues, and Y chromosome genes are implicated in sex differences in hypertension and cholesterol levels [2].

It has long been appreciated that gonadal sex steroids are crucial for sex differences. During gestation, ovaries develop unless the SRY gene is expressed, which leads to testes development. Activation of the hypothalamic-pituitary-gonadal (HPG) axis results in the production of kisspeptin in the hypothalamus which stimulates the release of gonadotropin-releasing hormone (GnRH). GnRH then regulates the production of follicle-stimulating hormone and luteinizing hormone, which stimulate the production of testosterone in the gonads which is converted to estrogen by the aromatase enzyme (Figure 1 A). Different levels of sex steroids lead to sexual development by differentially regulating gene expression throughout the body. The HPG axis is activated briefly during the neonatal period (mini-puberty) and then again at the start of puberty, when the majority of sexual maturation occurs [13].

**Figure 1.**
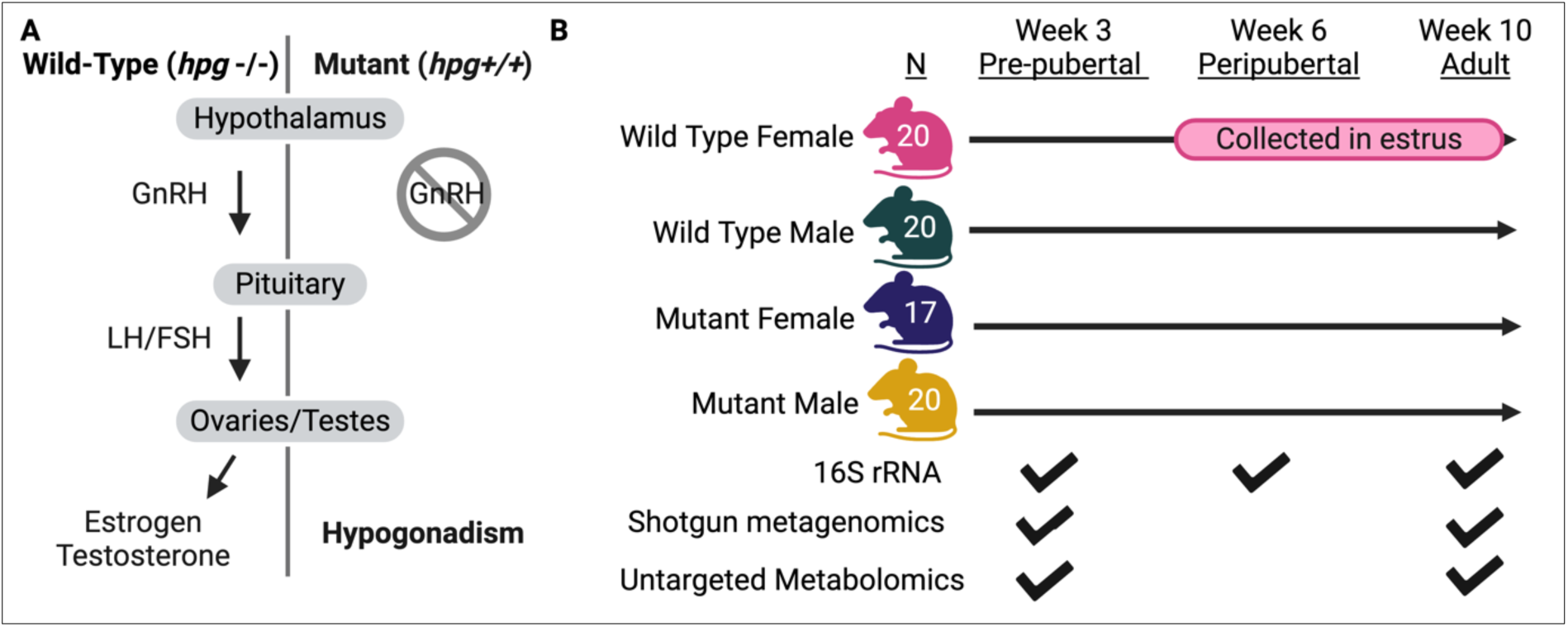
Experimental Design. (A) When the hypothalamic-pituitary-gonadal (HPG) axis is activated in wild-type mice (hpg-/-), the hypothalamus produces gonadotropin-releasing hormone (GnRH) that stimulates the anterior pituitary to secrete gonadotropins, luteinizing hormone (LH) and follicle-stimulating hormone (FSH). Gonadotropins regulate steroidogenesis and gametogenesis in the gonads (ovaries or teste), which results in secretion of sex steroids, sexual development, and reproductive competence. In mutant (hpg*+/+*) mice, the *Gnrh1* gene is truncated, so the GnRH peptide hormone is not produced, leading to an inactive HPG axis and hypogonadism. (B) Four experimental groups were included in the study: female and male wild-type mice and female and male mutant mice. N = mice per group. Fecal samples were collected at weeks 3 (pre-pubertal), 6 (peripubertal), and 10 (post-pubertal). For wild-type mice, fecal samples were only collected in the estrus phase of the estrous cycle in weeks 6 and 10. 16S rRNA amplicon sequencing was conducted on fecal samples at all weeks, and shotgun metagenomic sequencing and untargeted metabolomics were performed on fecal samples at weeks 3 and 10. Created with BioRender.com.

Like the mammalian host, the gut microbiome also undergoes sexual maturation. Understanding how this process occur is important because the gut microbiome is linked to many diseases that have a sex bias or perturbation of sex steroid levels, including inflammatory bowel syndrome [14], polycystic ovary syndrome [15], and type I diabetes [16]. Prior to puberty, studies observed no sex differences in the gut microbiome in either humans or mouse models [17–19], though studies comparing sex differences in the pre-pubescent gut microbiome are few. During puberty, sex differences in the composition of the gut microbiome develop and are maintained in reproductive-aged adults [20–23]. The cause of this sexual differentiation is not well understood, though it has been shown that gonadectomy and sex steroid replacement can shift gut microbiome composition in mice to resemble the microbiome of intact mice [24–27]. Since the timing of sexual differentiation of gut microbiome mirrors sexual development of the host, this provided a rationale to investigate if the HPG axis is required for sex differences in the gut microbiome and if so, when was this effect exerted e.g. during mini-puberty or puberty.

To understand how the HPG axis in the host influences maturation of gut microbial taxonomic composition and function, we investigated changes in the gut microbiome using feces as a proxy before and after puberty using a hypogonadal (hpg) mouse model (Figure 1 A). In this model, the HPG axis is not activated in homozygous mutant mice (hpg+/+) but is activated in wild-type mice (hpg -/-). By comparing how the microbiome developed in mutant and wild-type male and female mice, we showed that the HPG axis exerts a strong effect on the taxonomic composition of the gut microbiome during puberty. We also showed that HPG axis activation caused sex differences in bacterial species network structure and connectivity.

Moreover, the HPG axis was required for sex differences in microbial gene beta diversity, and microbial lipid metabolism. However, some sex differences in taxonomy and amine metabolism were also observed in mutant mice, indicating that an HPG-independent effect, e.g. sex chromosomes, may contribute to some sex differences in the gut microbiome.

## METHODS

### Hpg Mouse Model

Heterozygote hypogonadal (hpg) mice with a *Gnrh1^hpg^* mutation were purchased from Jackson Labs (RRID:IMSR_JAX:000804). The *Gnrh1^hpg^* mutation is a ∼33.5 kb deletion that removes two exons that encode most of the protein. Hpg heterozygous (hpg+/-^)^ male and female mice were crossed and the offspring were genotyped to identify hpg+/+ (mutant) or hpg-/- (wild-type) homozygous males and females. Two cohorts of breeding mice were generated with the aim of producing 20 mice per genotype and sex (Figure 1 B). Cohort A consisted of 8 female mutants, 10 female wild types, 11 male mutants, and 11 male wild types, while cohort B consisted of 9 female mutants, 10 female wild types, 9 male mutants, and 9 male wild types. In total, 77 wild-type and mutant mice were produced from 31 litters. Mice were housed in a vivarium with a 12-hour light (6:00 AM to 6:00 PM) and 12-hour dark cycle and had access to water and food (Teklad Global 18% Protein Extruded Diet; Envigo, Indianapolis, IN) *ad libitum*. After weaning at 21 days of age, mice were single housed to prevent coprophagy from influencing the gut microbiome. The University of California, San Diego Institutional Animal Care and Use Committee approved all animal procedures used in this study (Protocol Number S14011).

### Sample Collection and DNA Isolation

Fecal samples were collected and mice were weighed at 3, 6, and 10 weeks of age (Figure 1 B). Samples were frozen immediately after collection and stored at −80°C. To minimize potential variability induced by the estrous cycle, female wild-type samples were collected during estrus (determined by vaginal epithelial cell smears) at 6 and 10 weeks of age. Bacterial DNA was extracted from fecal samples and from two negative extraction controls and two ZymoBIOMICS Microbial Community Standard positive extraction controls (D6300 Zymo Research) with the DNeasy PowerSoil Pro Kit (Qiagen) according to the manufacturer’s instructions, and stored at −80°C.

### 16S rRNA Amplicon Sequencing

The 16S rRNA gene in all fecal samples and controls was PCR amplified using “universal” bacterial primers 515F and 806R for the V4 hypervariable region, with the 806R primers barcoded with unique 12-base pair (bp) Golay barcodes [28]. 16S rRNA amplification was also performed on two negative extraction controls, two negative PCR controls, two DNA extraction positive controls (ZymoBIOMICS Microbial Community Standard), and two PCR positive controls (ZymoBIOMICS Microbial Community DNA Standard).

Amplification was performed using the following steps: an initial denaturation temperature of 94°C for 3 minutes, then 25 cycles of 45 seconds denaturation, 60 seconds of 50°C annealing, 90 seconds of 72°C extension, then a 72°C final extension for ten minutes.

Amplicon sequencing libraries were prepared for the 231 experimental samples and 8 controls by The Scripps Research Institute Next Generation Sequencing Core. Amplicon products were cleaned with Zymo DNA Clean & Concentrator™-25 columns, quantified using a Qubit Flourometer (Life Technologies), and pooled. Sequencing libraries were prepared with the recommended Illumina protocol involving end-repair, A-tailing and adapter ligation. After library prep, libraries were PCR amplified with HiFi Polymerase (Kapa Biosystems) for 12 cycles. Quantitation, denaturation in 0.1 N NaOH, then dilution to 5pM of libraries preceded loading libraries onto an Illumina single read flow-cell for sequencing on the Illumina MiSeq.

### 16S rRNA sequence quality control and QIIME2

Forward read raw sequences were processed with QIIME 2 (version 2022.8) [29]. Reads and metadata were imported with qiime tools import. Reads were demultiplexed using QIIME 2 cutadapt [30]. This resulted in 8.58 million total reads with 34,616 average reads per sample. Sequence variants (SVs) were determined with dada2, truncating at 240 base pairs based on quality scores [31]. QIIME 2’s feature classifier plugin was used to assign taxonomy to SVs using a pretrained naïve Bayes classifier trained on reference database Silva 138 [32, 33]. Negative sequencing and DNA extraction controls and Zymo positive community sequencing and DNA extraction controls were evaluated for expected and unexpected sequences. Positive control community composition closely matched the expected genera abundances, while unexpected sequences in the positive controls were less than 1% of SVs. Negative controls had low sequence counts, with 18 to 880 times fewer counts than the lowest-count positive control. The composition of the negative controls greatly differed from the experimental samples. Sequences classified as mitochondria were removed with qiime taxa filter-table, leaving 741 SVs in the feature table. To deal with sparse features, SVs not present in at least five samples were removed using CurvCut [34], resulting in a final count of 333 SVs.

### Shotgun Metagenomic Sequencing

Shotgun metagenomic sequencing was performed on cohort B week 3 and 10 fecal sample DNA by the UCSD Institute for Genomic Medicine Genomics Center. Libraries were prepared by fragmenting 500 ng of genomic DNA from each sample by sonicating with Adaptive Focused Acoustics (E220 Focused Ultrasonicator, Covaris, Woburn, Massachusetts). This produced an average fragment size of 500 base pairs. The KAPA Hyper Prep Kit (KAPA Biosystems, Wilmington, MA, USA) was used to generate sequencing libraries following manufacturer’s instructions using 4 cycles of amplification. Library quality was assessed using High Sensitivity D1000 kit on a 4200 TapeStation instrument (Agilent Technologies, Santa Clara, CA, USA). The NovaSeq Sequencing System (Illumina, San Diego, CA, USA) was used to generate 150 bp paired-end reads to obtain ∼35 million reads per sample. An average of 76.4 million reads per sample were produced.

### Shotgun Metagenomics Sequence Quality Control

Paired-end reads were filtered and trimmed based on quality score and read length and adaptors and poly(G) tails were detected and removed by Fastp version 0.23.2 [35]. This decreased the average read length from 151 base pairs to 147 base pairs. The samples averaged 97 percent of bases with a Q20 or higher quality score, and 92 percent of bases with a Q30 or higher quality score. Quality-filtered and trimmed reads were then mapped to the Mus Musculus genome (Assembly GRCm39, Refseq GCF_000001635.27 downloaded from Ensembl) using Bowtie2 version 2.3.4.1[36]. Unmapped (non-host) reads were reconstructed to paired-end reads using SAMtools version 1.5 [37]. Singletons in forward and reverse read files for each samples were removed using fastq-pair version [38]. An average of 6.2 percent of reads were classified as host reads.

### Shotgun Metagenome Genome Assemblies and Annotation

Metagenome-assembled genomes (MAGs) were assembled with MEGAHIT version 1.2.9 [39]. Assembly was evaluated with QUAST version 4.4 [40]. Contigs over 250 bp were kept, resulting in 685,960 total contigs, of which 90,918 contigs were over 1,000 bp and 889 contigs were over 100,000 bp. Contigs were uploaded to KBase [41] and binned with MaxBin 2.0 v2.2.4 [42]. MAG quality was evaluated with CheckM v1.0.18 on KBase [43]. GTDB-Tk version 2.4.0 was used to annotate bins to the species level with the Gene Taxonomy Database [44, 45]. Salmon was used to determine relative counts of bins, and counts were summed for bins of the same species [46]. Prodigal version 2.6.3 was used to predict coding sequences (genes) from contigs [47]. Eggnog mapper and the Eggnog database version 5 were used to annotate gene orthologous groups [48, 49]. Salmon was used to generate a count table of coding sequences (genes) for each sample [47]. Gene features not present in at least four samples were removed using CurvCut [34].

### Metabolomics Mass Spectrometry and Annotation

West Coast Metabolomics performed metabolomic profiling on cohort B week 3 and 10 fecal samples. Metabolites were extracted using methyl-tert-butyl ether extraction and prepared as described previously [50]. The top layer (methanol fraction) was used for the lipidomics, while the bottom layer was used for analysis of biogenic amines. After extraction, sample fractions were reconstituted via vortexing for ten seconds and sonicating for 5 minutes, then centrifuged for 2 minutes at 16,100 x g. Spectra for lipids were produced with UHPLC ESI QTOF MS/MS (Agilent 6530(pos) 6546(neg) Santa Clara, CA), and spectra for biogenic amines and other small metabolites were produced using HILIC-ESI QTOF MS/MS (Sciex 6600 Framingham, MA).

Metabolites were annotated with a West Coast Metabolomics in-house mzRT library and MS/MS spectral matching with NIST/MoNA (National Institute of Standards and Technology/MassBank of North America) libraries. 17.6% of lipid metabolites and 9.1% of amine metabolites were able to be matched to a known compound. For each platform, peak heights were normalized to internal standards to account for within-series drifts of instrument sensitivity by dividing the raw metabolite (i) peak height within a sample (j) by the sum of peak heights (iTIC) for all internal standards within a sample (j), and multiplying that value by the average of all internal standards for all samples:

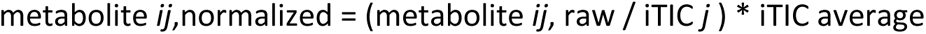

### Statistical Analyses

Statistical analyses and graphing were performed in R version 4.2.1. All graphs, with the exception of the network analyses, were generated using ggplot2 3.4.0. To compute family relative abundances, SVs were summed by family and family counts were made relative to the total sample counts. Family and gene feature tables were transformed using the centered-log-ratio (clr) for statistical analyses [51]. Linear mixed-effect model analysis on family clr-transformed counts was performed with R package nlme [52]. Euclidean distances of clr-transformed counts, Non-metric multidimensional scaling (NMDS) ordination of Euclidean distances, and PERMANOVA analysis of Euclidean distances were computed with Vegan [53]. Genera-level balances, coefficients, and AUC values were computed with coda4microbiome [54]. Random forest was performed on clr-transformed transformed metabolite abundances using a 70/30 train/test split using package randomForest [55]. For top important metabolites, normality of the data was determined with the Shapiro-Wilk test and t tests were performed on normalized peak heights. Within each analysis, p-values for multiple comparisons were corrected with the false discovery rate (FDR) method.

### Network Analyses

Network analysis was performed in python version 3.1 using matplotlib version 3.6.3 [56], network version 3.0 [57], numpy version 1.23.5 [58], pandas version 2.0.2 [59], and scipy version 1.13.0 [60]. 5000 bootstrapped replicates of the MAG species clr-transformed abundance dataset were created, where each bootstrap was randomly resampled with replacement to create new datasets. For each mouse group at weeks 3 and 10, Spearman correlations between species were calculated for each of the 5000 datasets. Networks were drawn and network statistics were calculated from the average of the species correlations on each of the 5000 datasets for edges with a correlation coefficient of 0.8 or higher. Network statistics were compared with binomial tests.

## RESULTS

### Broad taxonomic changes in murine gut microbiota are largely complete by 6 weeks of age

First, we profiled the family-level relative abundance of samples at each age using 16S data. Statistical analysis of the effect of age, hpg genotype, and sex on the ten most abundant bacterial families found that the relative abundance of these families was significantly different by sex, genotype, age, or a combination of these effects (Table 1). Overall family-level relative abundances at week 6 (peripubertal) were more similar to week 10 (adult) samples than week 3 (pre-pubertal) samples (Figure 1A). Compared to pre-pubertal samples, peripubertal and adult samples showed a greater relative abundance of *Muribaculaceae, Lactobacillaceae*, and *Oscillospiraceae*, and a diminished relative abundance of *Bacteroidaceae* and *Prevotellaceae* (Figure 1A). This pattern was consistent with beta diversity analyses at the SV level. NMDS of Euclidean distances showed that peripubertal and adult samples overlapped for all mouse experimental groups, while the pre-puberty cluster was distinct (Figure 1B).

**Table 1.** Results (p-values) of linear mixed-effect model analysis with top 10 most abundant bacterial families.

| Family | Sex | Genotype | Age | Sex:<br>Genotype | Sex:<br>Age | Genotype:<br>Age |
| --- | --- | --- | --- | --- | --- | --- |
| Lactobacillaceae | 0.8327 | 0.1529 | **2.50E-07** | 0.0901 | 0.2841 | **0.0092** |
| Lachnospiraceae | 0.9054 | 0.3094 | **5.43E-06** | **0.0011** | 0.0537 | **0.0032** |
| Bacteroidaceae | 0.3803 | 0.7363 | **8.29E-12** | **0.0021** | **0.0451** | **0.0011** |
| Muribaculaceae | 0.3803 | **0.0172** | **7.91E-13** | **3.0E-5** | 0.0599 | **3.04E-08** |
| Prevotellaceae | 0.9054 | 0.7363 | **0.00031** | 0.3311 | 0.5930 | **3.71E-06** |
| Oscillospiraceae | 0.8327 | 0.1529 | **7.10E-10** | **0.0130** | 0.0595 | **0.0104** |
| Deferribacteraceae | 0.9141 | 0.1577 | **0.0158** | **0.0101** | 0.6978 | 0.2595 |
| Ruminococcaceae | 0.9054 | 0.1529 | **0.04507** | **0.0136** | 0.1118 | 0.0944 |
| Rikenellaceae | 0.3803 | 0.4024 | **1.17E-09** | 0.9062 | 0.0595 | 0.9788 |
| Eggerthellaceae | 0.9141 | **0.0202** | **1.93E-12** | **0.0192** | 0.0595 | **0.0016** |

### HPG axis and sex influence family-level relative abundances

There was a strong effect of genotype on the relative abundances of 7 bacterial families, especially when age was taken into account (Table 1). There was a significant effect of genotype alone on *Muribaculaceae* and *Eggerthellaceae* abundances, while *Muribaculaceae, Eggerthellaceae, Lactobacillaceae, Lachnospiraceae, Bacteroidaceae, Prevotellaceae,* and *Oscillospiraceae* abundances were affected by a combination of age and genotype (Table 1). The combined effects of sex and genotype, or sex and age, also affected relative abundances of *Lachnospiraceae, Bacteroidaceae, Muribacilaceae, Oscillospiraceae, Deferribacteraceae, Ruminococcaceae,* and *Eggerthellaceae* (Table 1). *Bacteroidaceae* was also different by the combined effect of sex and age, and the FDR-corrected p values for this effect on *Lachnospiraceae, Muribaculaceae, Oscillospiraceae, Rikenellaceae*, and *Eggerthellaceae* were only slightly larger than a cutoff of 0.05, indicating that these families also may be affected by sex in an age-dependent manner.

Genotype was associated with clear differences in specific family-level bacterial abundances pre-versus post-puberty. *Prevotellaceae* abundances were higher in wild type mice pre-puberty, but not in adults. Members from the *Muribaculaceae* family were more abundant in mutant mice at peripuberty and adulthood, and *Lachnospiraceae* were more relatively abundant in wild type mice at peripuberty and adulthood (Figure 1A).

### Genera balances differentiate post-pubertal gut microbial communities by sex and genotype

To determine how age and reproductive development influence microbial community composition at a lower taxonomic level, we modeled microbial balances over time for each mouse group (wild-type female, wild-type male, mutant female, and mutant male). Genera coefficients indicate the relative proportion that the bacterial genus abundance contributed to the balance. The 16S dataset was used because the sample size was double that of the metagenomic dataset, and genera-level models were used because they had considerably higher AUC scores (better fit to the data) than SV-level models. Balance analysis using genera abundances differentiated microbial communities by age, genotype, and sex. With regards to age, *Candidatus arthromitus* and *Bacteroides* were identified as being more abundant in week 3 (pre-puberty) balances for all mouse groups compared to weeks 6 and 10 (Figure 3 A-D) while *Anaerotrunctus* was more abundant in week 3 balances for *hpg* mutants and female wild-type mice (Figure 3 A-C). For wild type, but not mutant, mice the week 3 genera coefficients also included *Alloprevotella* (Figure 3 C-D). While no genera were more abundant in week 10 balances across all four groups, some week 10 genera coefficients were conserved by genotype: *Alistipes, Lactobacillus,* and *Intestimonas* were more abundant in week 10 wild type balances and *Muribaculaceae* was more abundant in week 10 balances of mutant mice, regardless of sex (Figure 3 E).

**Figure 2.**
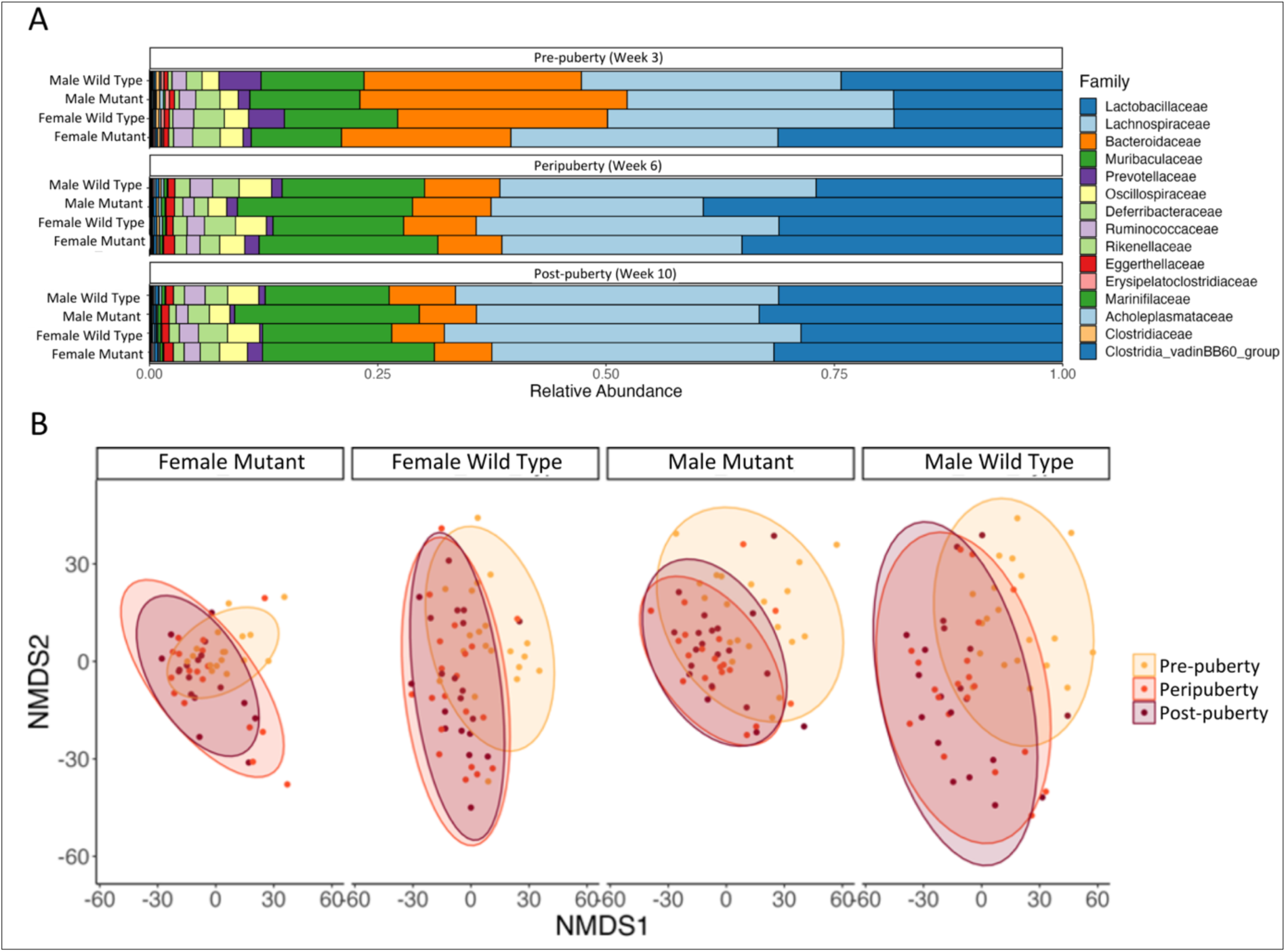
Age (week), hpg genotype, and sex influence gut microbial family abundances and beta diversity computed from 16S sequencing data. (A) Bacterial family relative abundances of each mouse group (male wild type, male mutant, female wild type, and female mutant mice) at age 3 (pre-puberty), 6 (peri-puberty), and 10 weeks (post-puberty). NMDS ordinations by mouse group for (B) Euclidean distances identified by different colors by age.

**Figure 3.**
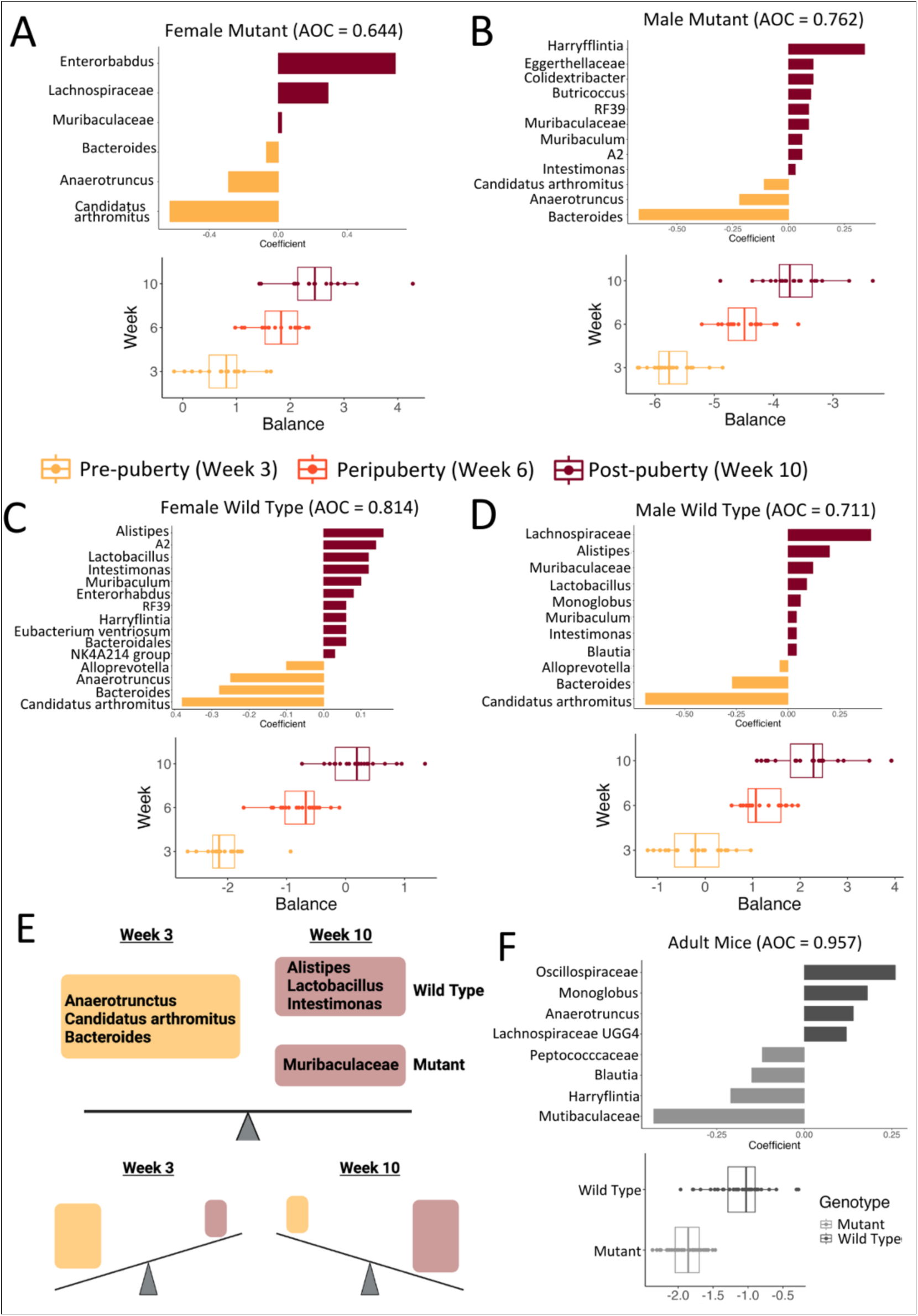
Age-based genera balances differentiate adult mice by sex and hpg genotype. Genera coefficient values and balance scores for balances defining age-based community differences (A) female mutant, (B) male mutant, (C) female wild-type, and (D) male wild-type mice 3, 6 and 10 weeks of age. (E) Summary schematic of common genera defining longitudinal balances at week 3 and at week 10 based on hpg genotype. (F) Genera balance coefficient values and balance distribution for mutant vs wild-type mice at week 10.

Week 10 genera coefficients also differed by sex for both wild-type and mutant mice, with *Entoerorhabdus* and a *Lachnospiraceae* genus more relatively abundant in adult mutant females, while *Harryflintia,* an *Eggerthellaceae* genus, *Colidextribacter, Butrycoccus, RF39, Muribaculum, Intestimonas, and A2* were more relatively abundant in adult mutant male (Figure 3 A-B)*. A2, Muribaculum, Enterorhabdus, RF39, Harryflintia, the Eubacterium ventriosium* group, and a *Bacteroidales* genus were more relatively abundant in adult female wild-type balances, while *Lachnospiraceae* genus, a *Muribaculaceae* genus, *Monoglobus,* and *Blautia* were more relatively abundant in the adult male wild-type balances (Figure 3 C-D).

Given that age-based balances for each group showed conserved differences between mutant and wild-type mice regardless of sex, we modeled balances differentiating mutant and wild-type communities for both sexes in adulthood at 10 weeks of age. The AUC of this model was higher than the age comparison models, indicating the strength of this method at discriminating samples based on genotype in adult mice. Similar to the results of the age-based balances, *Muribaculaceae* was more relatively abundant in mutant mice in adulthood, while *Alistipes* was more relatively abundant in wild-type mice. The conserved difference between wild-type and mutant mice, in both the age-based and adult *hpg*-based balances, indicates a strong effect of the HPG axis on development of the gut microbiome during the pubertal period.

### HPG axis activation affects sex differences in beta diversity at the species level

Since the 16S dataset did not annotate most SVs to the species level, we used metagenome-assembled genomes (MAGs) to identify bacterial species and determine how age and the HPG axis altered bacterial community composition. Species-level beta diversity analysis using Euclidean distances between samples identified significant effects of age, genotype, and sex, with age having the strongest effect followed by sex and genotype (Table 2). NMDS ordination of Euclidean distances on bacterial species showed that slight sex differences in beta diversity were apparent before puberty in both wild type and mutant mice (Figure 4A). Post-puberty, dispersion in mutant mice increased for both males and females compared to pre-puberty and sex differences in beta diversity were absent (Figure 4 A). For wild-type mice, adult male wild-type beta dispersion was much smaller compared to pre-pubertal samples, resulting in a sex difference in wild-type mice post-puberty (Figure 4 A).

**Figure 4.**
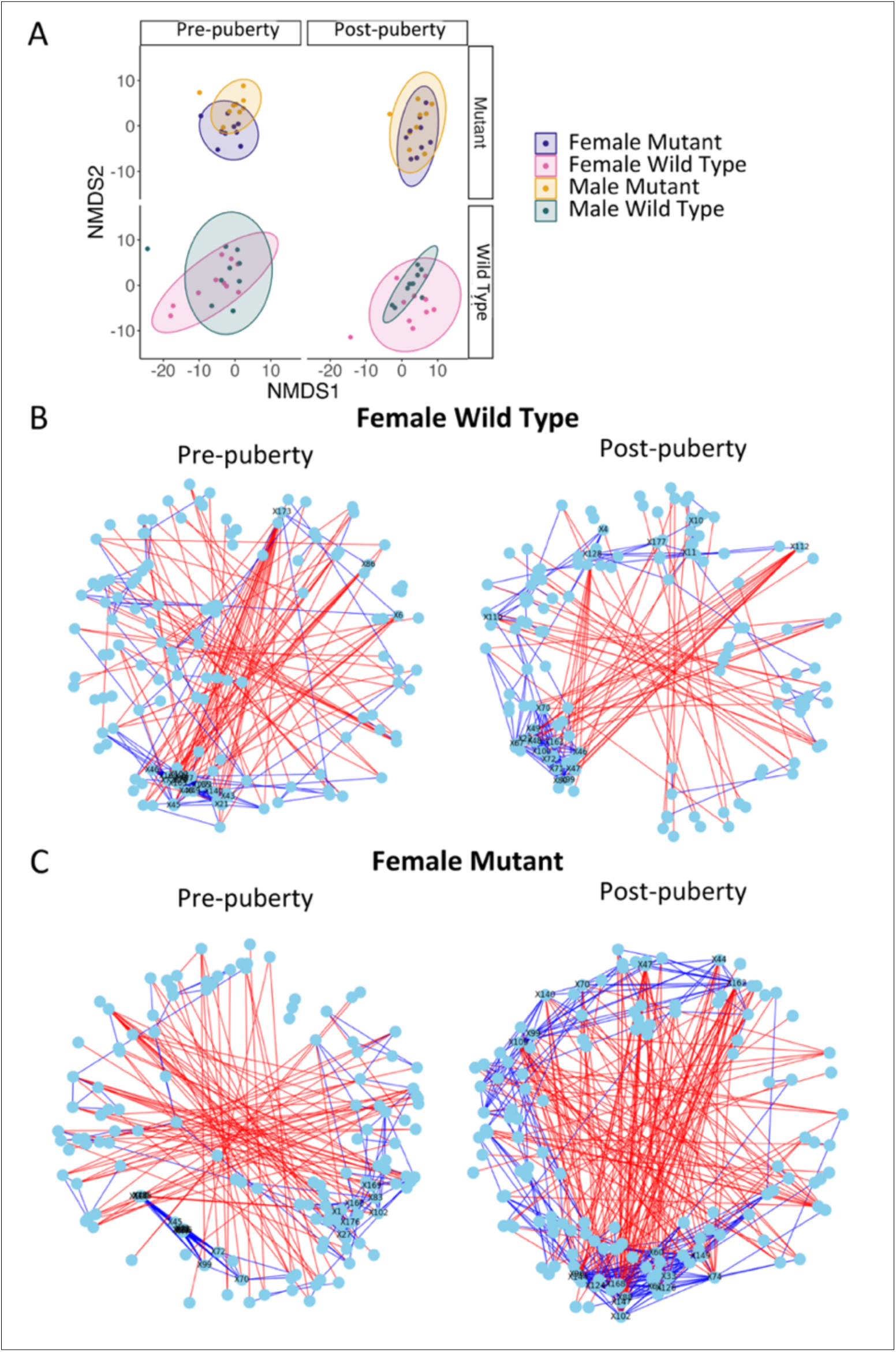
Puberty alters fecal bacterial community composition and networks at the species level in metagenomes. (A) NMDS ordination of Euclidian distances for species abundances between samples demonstrate sex differences before puberty (wild type and mutants) and after puberty (wild-type). Visualizations of pre-puberty and post-puberty for female wild-type (B) and female mutant (C) mice. Blue lines are positive correlations and red lines are negative correlations. Nodes are colored in light blue and represent species. The top 20 species with the most nodes are labeled with numbers corresponding to species listed in Supplemental Table 1.

**Table 2.** PERMANOVA results for Euclidean distances computed for MAG bacterial species. P-values less than 0.05 are bolded.

|  | Species |  |
| --- | --- | --- |
|  | R2 | Pr(>F) |
| **Sex** | 0.0336 | **0.0012** |
| **Genotype** | 0.0224 | **0.0280** |
| **Week** | 0.1013 | **0.0001** |
| **Sex:Genotype** | 0.0120 | 0.4283 |
| **Sex:Week** | 0.0103 | 0.6113 |
| **Genotype:Week** | 0.0106 | 0.5621 |
| **Sex:Genotype:Week** | 0.0084 | 0.8186 |

### HPG axis activation during puberty in females results in lower network connectivity

To further investigate how age and the HPG axis modulated community composition at the species level, we used a bootstrap approach (5000 replicates) to construct correlation networks based on MAG-derived bacterial species counts for each mouse group at weeks 3 and 10 (before and after puberty). Network structure and connectivity metrics were similar for all 4 groups of mice before puberty (Supplemental Figures 3.1 and 3.2). In contrast, many network metrics including number of edges (connections), degree centrality (connections per species), transitivity (degree to which nodes cluster together), and density (ratio of actual to potential edges) increased significantly from pre-to-post puberty with the exception of metrics for female wild-type mice which decreased (Table 3, Supplemental Figure 1). As visualized, a reduction in edges also occurred in wild-type females post-puberty compared to pre-puberty (Figure 4 B), which contrasted sharply with the increase in number of edges that occurred in other groups including female mutant mice (Figure 4 C).

**Table 3.**
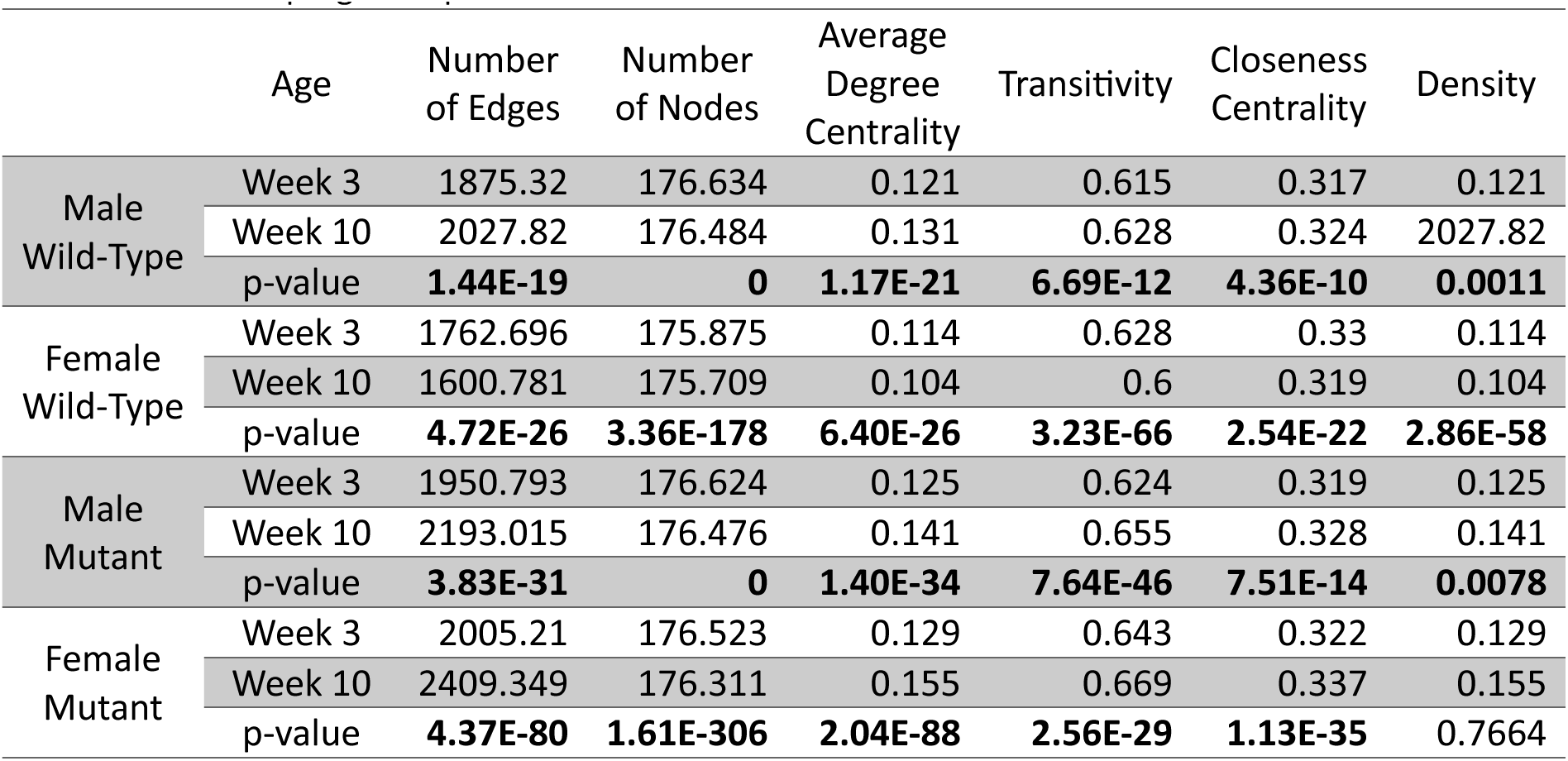
MAG species network statistics for week 3 and week 10 (5000 bootstrap replicates) and Binomial tests p-values comparing week 3 and week 10 networks. P values are FDR corrected. P values listed as 0 are not actually zero, but lower than the program’s precision. P values < 0.05 are bolded.

While the overall network structure was visually similar in all 4 groups before puberty (Supplemental Figure 2), the structure of female wild-type network differed from the other networks after puberty. Additionally, in adults, the mutant networks appeared strikingly different to wild-type networks, with more overall edges (Figure 3-5A-D, Table 3), and all adult networks except the wild-type female network contained two distinct clusters of positively correlated species (Figure 5A-D).

**Figure 5.**
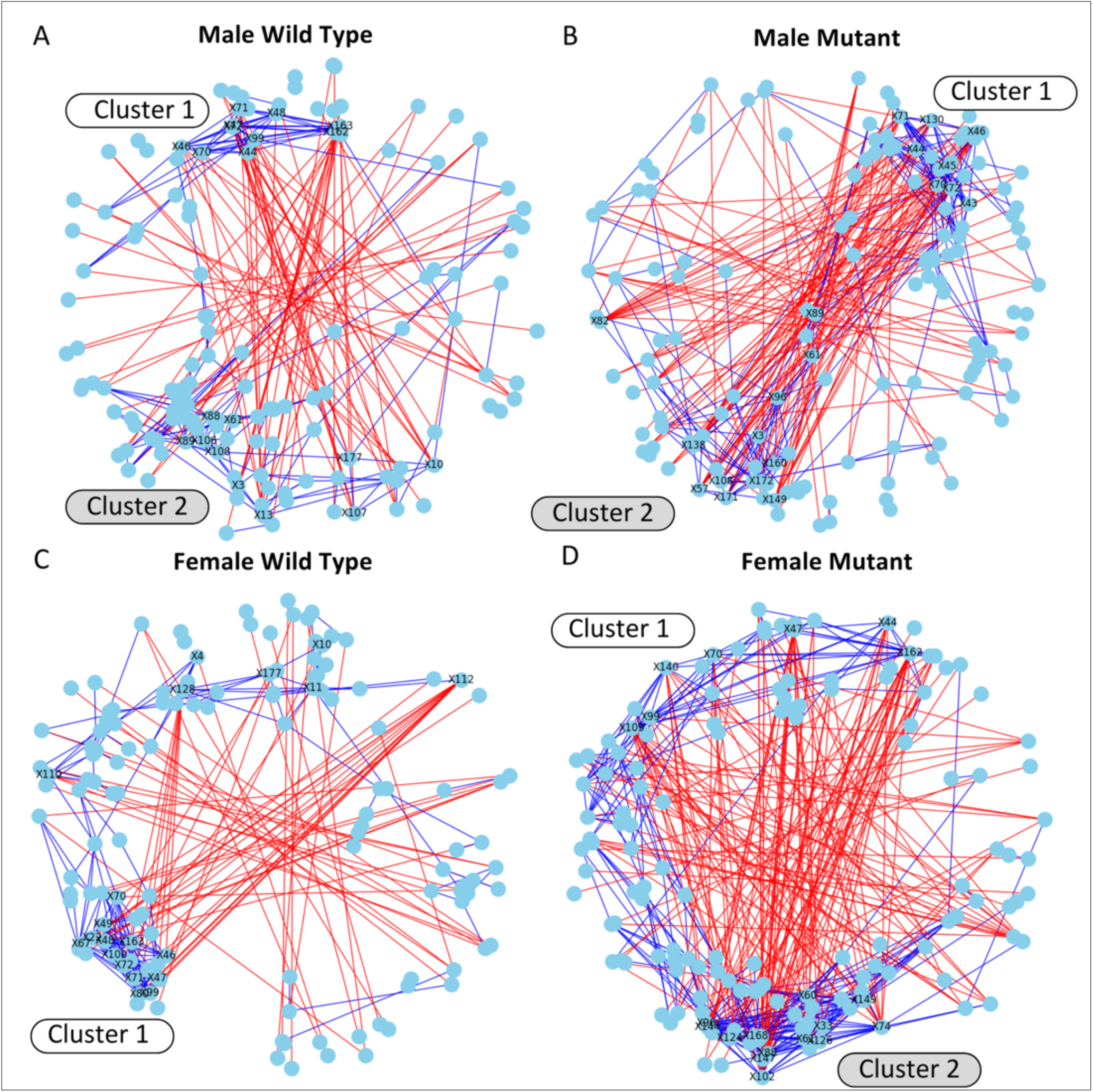
HPG axis influences structure of adult fecal bacterial networks. Visualization of week 10 networks for (A) male wild-type, (B) male mutant, (C) female wild-type, and (D) female mutant mice. Blue lines indicate positive correlations and red lines negative correlations. Nodes are colored in light blue and represent species. The 20 most connected nodes in each network are labeled with numbers corresponding to species listed in Supplemental Table 1. Cluster 1 consists primarily of *Muribaculaceae* species while Cluster 2 consists primarily of *Lachnospiraceae* species.

### Pre-pubertal networks display HPG-independent sex differences in clustering patterns

To investigate how age, HPG axis, and sex influenced which species were most connected in each network, we determined the twenty-most connected nodes in each network. The majority of these species in both pre- and post-pubertal networks belonged to *Muribaculaceae* and *Lachnospira*ceae families (Supplemental Tables 3.2 and 3.3). The specific species that were most connected varied between networks, though members of the *Duncaniella* and *Paramuribaculum* genera (family *Muribaculaceae*) were among the most connected nodes across networks (Supplemental Tables 3.2 and 3.3).

While overall network metrics were similar between the 4 groups of mice before puberty, we did observe network structure-related sex differences in both wild-type and mutant mice with regards to highly connected bacterial species. In pre-pubertal communities, highly connected nodes in males were dispersed through the networks and did not display strong clustering, though two nodes, an unclassified bin (labeled 168) and an unidentified *Lachnospiraceae* species UBA3282 (labeled 154), clustered together in both networks (Figure 4 D, E). Prepubertal female networks showed stronger clustering of networks than in males, with most of the highly connected nodes clustering together, containing species predominantly in *Muribaculaceae*, and *Lachnospiraceae* (Figure 4 F, G).

### Sex-specific species clustering in adult networks is driven by HPG axis activation in females

In adult wild-type networks, clustering of species was sex-specific in wild-type mice but not mutant mice, indicating an HPG-dependent sex difference that developed during puberty (Figure 5). In adults, most highly connected species fell into positively connected clusters that were dominated by specific families; one cluster type consisted primarily of *Muribaculaceae* species (Cluster 1) and the other cluster type consisted primarily of *Lachnospiraceae* species (Cluster 2) (Figure 5). The female wild-type network was the only adult network that lacked Cluster 2, and the only adult network where *Eggerthellaceae* species were also part of Cluster 1. Comparison of the highly connected species among the adult networks found that *Evtepia* species (labeled 88 and 89, unclassified *Eubacteriales*) and *Coprobacter secundus* (labeled 61, family *Barnesiellaceae*), in Cluster 2, were among the most-connected nodes for all networks with the notable exception of wild-type females. In contrast, only the female wild-type network included *Mucispirillum schaedleri* (labeled 128, family *Muribaculaceae*) and *Lawsonibacter* species (labeled 112, family *Oscillospiraceae*), which were not in a cluster (Figure 5C).

### Activation of the HPG axis during puberty leads to greater beta diversity of bacterial genes

After determining how sex and genotype influenced microbial taxonomic composition, we then investigated how putative bacterial functions differed before and after puberty. We examined differences in the overall set of predicted bacterial genes and orthologous groups of genes. PERMANOVA tests on Euclidean distances of gene and ortholog abundances between samples revealed a significant effect of sex, genotype, and age (pre-vs. post-puberty) on beta diversity, with the strongest effect due to age (Table 3). The effect of genotype was distinct for microbial genes and orthologs. For genes, the beta-diversity of wild-type and mutant samples in males and females was similar before puberty, while after puberty, wild-type samples had greater dispersion than mutant samples, reflecting an effect of activation of the HPG axis (Figure 6 A). In contrast, beta dispersion for all groups was reduced post-puberty for orthologs (Figure 6 A), indicating that the HPG axis axis effect on bacterial genes is due to changes in the abundance of specific genes instead of gene categories. Sex had less of an effect on the beta diversity for genes and orthologs (Figure 6 B).

**Figure 6.**
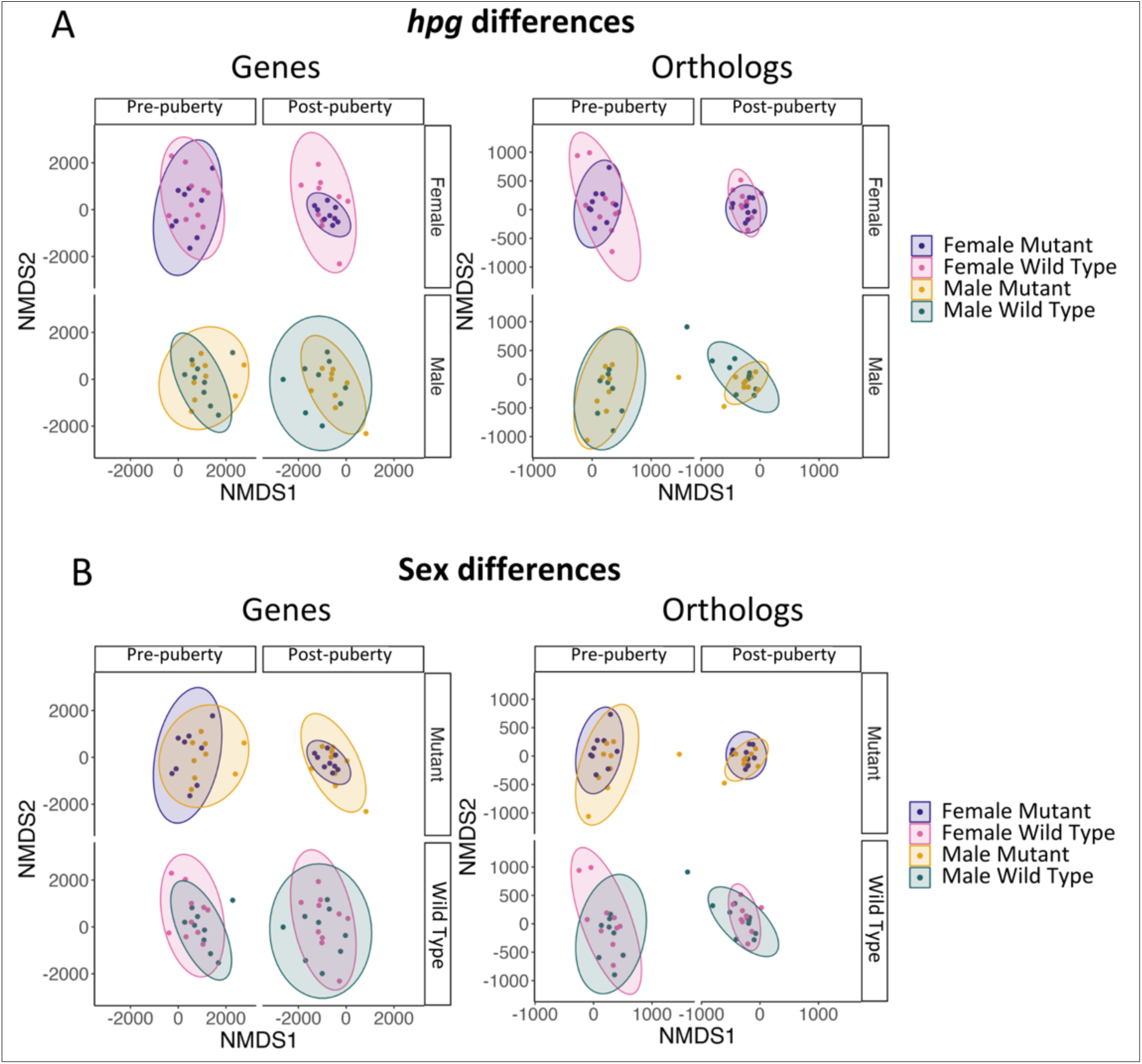
Microbial genes and orthologous groups differ by age, genotype and sex. NMDS ordination of Euclidian distances comparing (A) genotype and (B) sex differences for genes and ortholog abundances between samples.

### Activation of HPG axis during puberty leads to sex differences in lipid metabolites

Since we observed a strong effect of the HPG axis on the beta diversity of bacterial genes in mice after puberty, we then investigated the influence of age, sex and genotype on metabolites involved in lipid metabolism. PERMANOVA analysis indicated that beta diversity of lipid metabolites was different by age (pre-vs post-puberty) and by the combined effect of sex, genotype, and age (Table 5). NMDS ordination indicated that for both mutant and wild-type mice, sex differences were not apparent before puberty, but sex differences emerged in wild-type mice after puberty (Figure 7 A).

**Figure 7.**
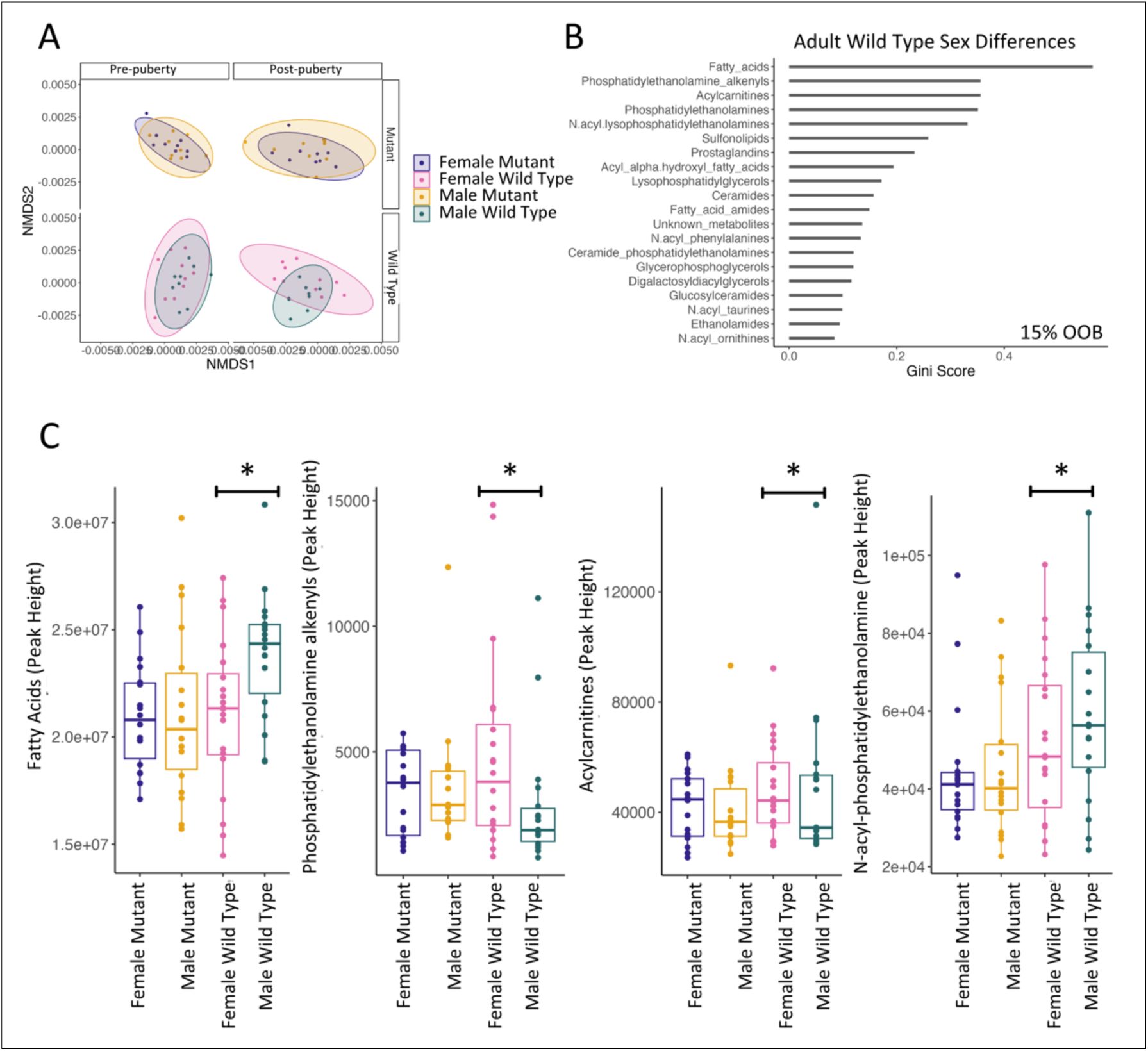
Gut lipid metabolites differ by age, sex, and genotype. (A) NMDS ordination of Euclidian distances showing sex differences within each genotype pre- and post-puberty. (B) Gini score for the most important lipid classes differentiating male and female wild-type mice. (C) Abundances of lipid classes that were significantly different by sex. OOB stands for out-of-bag error.

**Table 4.** PERMANOVA results for Euclidean distances computed for Gene and Ortholog abundances. P-values less than 0.05 are bolded.

|  | Genes |  | Orthologs |  |
| --- | --- | --- | --- | --- |
|  | R2 | P | R2 | P |
| **Sex** | 0.0210 | **0.0012** | 0.0222 | **0.0015** |
| **Genotype** | 0.0230 | **0.0002** | 0.0226 | **0.0010** |
| **Age** | 0.0554 | **0.0001** | 0.0671 | **0.0001** |
| **Sex:Genotype** | 0.0159 | 0.0592 | 0.0157 | 0.0781 |
| **Sex:Age** | 0.0110 | 0.8736 | 0.0109 | 0.8396 |
| **Genotype:Age** | 0.0143 | 0.1838 | 0.0141 | 0.1973 |
| **Sex:Genotype:Age** | 0.0103 | 0.9664 | 0.0098 | 0.9750 |

**Table 5.** PERMANOVA results for Euclidean distances computed for metabolite datasets. P-values less than 0.05 are bolded.

|  | Lipids |  | Amines |  |
| --- | --- | --- | --- | --- |
|  | R2 | P | R2 | P |
| **Sex** | 0.02128 | 0.0982 | 0.02698 | **0.0436** |
| **Genotype** | 0.01287 | 0.3129 | 0.01104 | 0.4527 |
| **Age** | 0.12143 | **0.0001** | 0.05328 | **0.0014** |
| **Sex:Genotype** | 0.01768 | 0.1637 | 0.02444 | 0.0650 |
| **Sex:Age** | 0.01501 | 0.2409 | 0.04057 | **0.0078** |
| **Genotype:Age** | 0.01585 | 0.2166 | 0.01039 | 0.4862 |
| **Sex:Genotype:Age** | 0.04389 | **0.0045** | 0.03033 | **0.0256** |

We also used Random Forest to determine which lipid classes were predictive in classifying sex differences before or after puberty (Figure 7 B). Interestingly, the model for adult wild-type mice had a significantly lower out-of-bag (OOB) error (15%) indicating a better fit to the data, while the other models had OOB errors of <30% (Table 6). For the 5 most important lipid classes differentiating wild-type males and females, all but phosphatidylethanolamines were significantly different by sex for adult wild-type mice, and none were significantly different by sex for adult mutant mice (Table 7). Fatty acids and N-acyl-lyphophosphatidylethanolamines were more abundant in wild-type males than females, and phosphatidylethanolamine alkenyls and acylcarnitines (fatty acid metabolites) were more abundant in wild-type females than males (Figure 7C). Taken together, these results indicate that activation of HPG axis during puberty drives sexual differentiation of gut lipid metabolism.

**Table 6.** Out-of bag error and accuracy for Random Forest models on sex differences for pre-pubertal (week 3) or post-pubertal (week 10) wild-type or mutant mice.

| Dataset | Age | Week/Genotype | Out-of-bag error | Accuracy on test set |
| --- | --- | --- | --- | --- |
| Amines | Pre-Puberty | Mutants | 42% | 42% |
| Amines | Post-Puberty | Mutants | 25% | 25% |
| Amines | Pre-Puberty | Wild Type | 39% | 33% |
| Amines | Post-Puberty | Wild Type | 30% | 50% |
| Lipids | Pre-Puberty | Mutants | 33% | 67% |
| Lipids | Post-Puberty | Mutants | 50% | 50% |
| Lipids | Pre-Puberty | Wild Type | 54% | 16% |
| Lipids | Post-Puberty | Wild Type | 15% | 67% |

**Table 7.** T-test results (p-values) for lipid class sex differences in adult mice within each genotype. P-values are FDR-corrected with values < 0.05 in boldface.

|  | Wild Type | Mutant |
| --- | --- | --- |
| Acylcarnitines | **0.0257** | 0.8636 |
| Fatty acids | **0.0044** | 0.3539 |
| N-acyl-lysophosphatidylethanolamines | **0.0360** | 0.5963 |
| Phosphatidylethanolamine alkenyls | **0.0257** | 0.5965 |
| Phosphatidylethanolamines | 0.1417 | 0.3606 |

### Sex differences in amine metabolites are driven by HPG-dependent and -independent effects

In addition to age, PERMANOVA analysis of fecal amine metabolites showed that beta diversity was significantly different by sex, the combined effect of sex and age, and the combined effect of sex, genotype, and age (Table 5). NMDS ordination showed that, while slight sex differences in amine beta diversity occurred before puberty, sex differences were readily apparent after puberty in both wild-type and mutant mice (Figure 8 A).

**Figure 8.**
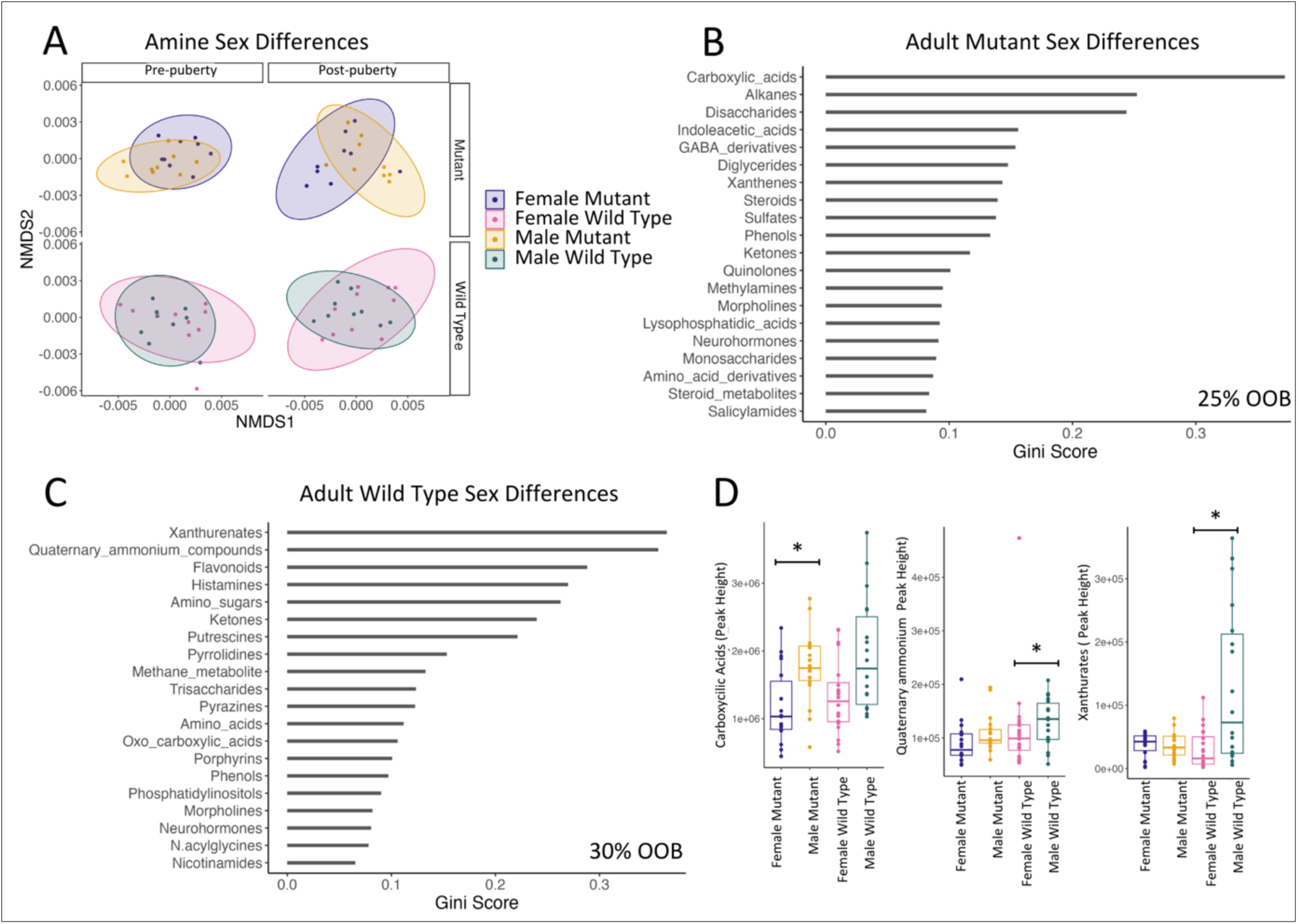
Gut lipid metabolites differ by age, sex, and genotype. (A) NMDS ordination of Euclidian distances showing sex differences within each genotype pre- and post-puberty. (B) Gini score for the most important lipid classes differentiating male and female wild-type mice. (C) Abundances of lipid classes that were significantly different by sex. OOB stands for out-of-bag error.

To determine which amines differentiated mice by sex before and after puberty, we employed Random Forest for both wild-type and mutant mice. For pre-pubertal mice, model out-of-bag (OOB) error was high while for post-pubertal mice, the model OOB error was lower but model accuracy on the test set was quite low (25% and 50%, respectively; Table 6). To investigate whether the levels of metabolites in specific amine classes identified in the Random Forest models as differentiating adult males and females were different between the sexes, we compared the abundances of the top 5 most-important amine classes for each model and found that three of the top amine classes were significantly different between males and females (Table 8, Figure 8 B-C). Interestingly, carboxylic acids were different between mutant males and females, while quaternary ammonium compounds and xanthurenates were different between wild-type males and females (Figure 8 D). These results indicate that adult sex differences in gut amine metabolites are due to both HPG-dependent and -independent effects.

**Table 8.** FDR corrected t-test results (p-values) for amine class sex differences in adult mice within each genotype. P-values <0.05 are in boldface.

|  | Wild-type Mice | Mutant Mice |
| --- | --- | --- |
| Alkanes | 0.1204 | 0.9811 |
| Amino sugars | 0.0525 | 0.7692 |
| Carboxylic acids | 0.0525 | **0.0345** |
| Disaccharides | 0.2168 | 0.2712 |
| Flavonoids | 0.0525 | 0.1752 |
| GABA derivatives | 0.6987 | 0.5274 |
| Histamines | 0.0525 | 0.5274 |
| Indoleacetic acids | 0.1057 | 0.1680 |
| Quaternary ammonium compounds | **0.0085** | 0.2002 |
| Xanthurenates | **0.0119** | 0.2712 |

## DISCUSSION

Multi-omics analysis of the fecal microbiome using a hypogonadal mouse model revealed that activation of the HPG axis during puberty was required for sexual differentiation of the gut microbiome including bacterial composition, putative gene functions, and lipid metabolism. This study also provided evidence sex-differentiated maturation of the gut microbiome outside of the HPG axis. In addition to determining a strong effect of age on the gut microbiome, sex differences in the gut microbiome were detected in both wild-type and mutant mice before puberty. This pre-pubertal sex-associated differentiation indicates that sex chromosome genes may also influence specific sex differences in gut microbial taxa and function.

### Age-dependent maturation of the gut microbiome during puberty

Comparisons of microbial diversity before, during, and after the pubertal window between mice with and without a functioning reproductive axis allowed us to discriminate the effects of sexual development on the gut microbiome from those due to other developmental factors. Analysis of gut microbiota taxa and putative functions showed that age, independent of HPG axis activation, strongly shaped the gut microbiome during the pubertal period. This finding suggests that the common notion in the field that sexual development is the primary factor driving maturation of the gut microbiome during puberty [61] is incomplete. It also contradicts previous hypotheses that age-related gut microbiome maturation is complete before puberty [62, 63]. Our data showed that age, independent of sex or HPG axis activation exerts a considerable effect on broad changes in the gut microbiota composition and putative functions. Additionally, we identified a common set of bacteria that differentiated pre-pubertal mice from adult mice in all four groups. This indicates that factors independent of sexual development, such as weaning and growth, result in reduced abundance of these genera in adult mice.

Analysis of orthologous groups of genes annotated from MAGs also indicated an HPG-independent effect of age during puberty: the beta dispersion of orthologous groups in all four mouse groups collapsed post-puberty. Orthologs, groups of genes that have evolved from a common ancestor, represent a higher functional level than genes. Thus, the substantial reduction of beta dispersion we observed suggests that, as the mice mature throughout puberty, the broad functions of the gut microbiota converge to a shared functional profile and that this process is not affected by activation of the HPG axis.

### HPG axis activationis a critical factor in gut microbiome maturation

Analysis of taxonomic diversity, particularly at the level of bacterial families, showed that the HPG axis was required for full maturation of the gut microbiome during puberty. It also showed the impacts of the HPG axis was similar in both male and female mice. The two families that were most affected by the HPG axis were the *Muribaculaceae* and the *Lachnospiraceae*. Members of the *Muribaculaceae* family were significantly more abundant in adult mutant mice compared to wild-type mice. Bacteria in this family are known to degrade complex carbohydrate, including host mucins, and they are also know to contain genes for bile acid transformation [64]. Previous work showed that *Muribaculaceae* species were especially abundant in mutant adult mice in microbial communities of the small and large intestine [65]. Members of the *Lachnospiraceae*, on the other hand, were more abundant in wild-type mice at peripuberty and adulthood compared to mutant mice, a result supported by a balance analysis that showed one specific genus of *Lachnospiraceae* to be more abundant in wild-type mice than mutant mice. Our previous study also found higher abundances of *Lachnospiraceae* in the cecum lumen of wild-type mice compared to mutant mice[65]. Studies have shown that *Lachnospiraceae* degrade host mucins and produce short-chain fatty acids that can affect host inflammation [66, 67]. Taken together, these results indicate that HPG axis activation during puberty suppresses abundance of bacteria in the *Muribaculaceae* family and promotes abundance of *Lachnospiraceae*.

Analysis of genes annotated from MAGs indicated that the HPG axis is also necessary for diversification of bacterial community gene functional diversity in adults. Before puberty, all four groups of mice showed high beta dispersion in gene abundance, but post-puberty, beta dispersion collapsed in mutants while it increased in wild-type mice, indicating that activation of the HPG axis generates higher levels of inter-sample genetic differentiation in the gut microbiome.

### HPG axis is required for sex differentiation of the gut microbiome during puberty

Analysis of gut microbial diversity between the sexes in both wild-type and mutant showed that HPG axis activation during puberty led to substantial sex differences in the gut microbiome, particularly in the comparisons of species-level beta diversity and correlation networks. In wild-type mice, sex differences in species beta dispersion developed during the pubertal period, while this was not true for mutant mice.

Species-level bacterial community networks revealed that HPG-dependent sex differences in adult wild-type species networks were absent before puberty. However, post-puberty, the female wild-type microbial network was much sparser than the other groups and all its connectivity metrics were substantially lower. Previous studies in non-gut microbial ecosystems have shown that environmental disruption destabilize communitiey networks, leading to sparser networks and lower levels of overall connectivity, similar to what we observed in the adult female wild-type mouse networks [68]. It is possible that activation of the HPG axis in female mice during puberty results in greater physiological change in the gut of females (i.e., greater environmental instability) than in the other groups, which is reflected in the networks. One possible reason for this greater level of instability in females might be changes in the intestinal environment due to cyclic changes in hormone levels during estrous. Variability in gastrointestinal transit time provides another potential explanation. Transit time has been shown to be highly responsive to estrogen and progesterone levels and has been identified as an important variable in sex differences within the human gut microbiome [69, 70].

A closer look at the networks found two distinct species clusters in the adult mice that were negatively correlated with one another. Cluster 1, was dominated by *Muribaculaceae* species, while Cluster 2 was dominated by members of the *Lachnospiraceae.* There out of the four mouse groups had both clusters, but Cluster 2 was missing from female wild-type mice. While direct interactions between connected species cannot be assumed from co-occurrence networks, organization of species networks into co-occurrence clusters grouped at the family level likely indicates that the species in these clusters occupy similar niches within the gut environment or are mutually dependent and that different clusters occupy different niches [71]. Network clustering into groups based on higher-level taxonomy has been seen in mouse models of Alzheimer’s disease and exercise stress [72, 73]. While *Muribaculaceae* and *Lachnospiraceae* are known to degrade carbohydrates in the gut [67, 74], *Lachnospiraceae* species are key butyrate producers and have also been shown to ferment amino acids [75–77]. That the adult wild-type female species networks were missing the *Lachnospiraceae* co-occurrence cluster suggests that the species occupying this niche were not selected by activation of the HPG axis in female mice.

Additionally, activation of the HPG axis during puberty was required for sexual differentiation of lipid metabolite beta diversity. We note that all of these HPG-dependent sex differences in microbial taxa and functions were absent pre-puberty, which ruled out the neonatal period (mini-puberty) as a contributing factor to sexual differentiation of the gut microbiome via activation of the HPG axis.

### Sex chromosome genes influence some sex differences in the gut microbiome

This study also showed that some sex differences in the gut microbiome were independent of the HPG axis, indicating that gene dosage effects of sex chromosomes likely play a role in sexual differentiation of some aspects of the gut microbiome. Specifically, we found sex differences in beta diversity in pre-pubertal mutant mice that diminished post-puberty, indicating that sex chromosome effects may be transient or masked in adulthood by other factors. While studies on pre-pubertal sex differences are few [19], some studies have shown sex differences in pre-pubescent mice in gene expression in the liver and intestine [78, 79].

Additionally, metabolomic analyses indicated that adult sex differences in fecal amine metabolites were also HPG independent. Beta diversity differences in amines were most evident in adult mutant mice, indicating that sex differences resulting from sex chromosome genes were largely masked by activation of the HPG axis during puberty in wild-type mice (Figure 8). A similar masking phenomenon wherein sex steroids were antagonistic to sex chromosome effects has been previously described in a mouse model with a sexually-dimorphic immune response [80]. For specific amine classes, there were both HPG-dependent and independent sex effects. Carboxylic acids differentiated male and female mutant mice, while quaternary ammonium compounds and xanthurates differentiated male and female wild-type mice. Altogether, these results reveal complex interactions between gonadal sex steroids and sex chromosomes in shaping functional sex differences in the gut microbiome.

### Potential mechanisms for sex-specific maturation of the gut microbiome

The identification of HPG-independent effects on maturation of the gut microbiome is a novel finding and has not previously been proposed in the field despite previous studies that demonstrated gene dosage effects of sex chromosomes on host physiology. Given the abundance of immune genes on the X chromosome, immune action in the gut is a likely candidate for sexual differentiation of the gut microbiome due to sex chromosomes. Indeed, differential abundance of autoimmune genes in the small and large intestine have been observed between pre-pubertal female and male mice [79]. Another mechanism through which this could occur is differential expression of X/Y homologous genes: many of these genes exhibited differential expression between adult females and males in the liver, colon, and small intestine of mice [12].

However, given that most sex differences identified in study were dependent on activation of the HPG axis, most of the sexual maturation of the gut microbiome is likely due to direct or indirect action of gonadal sex steroids. The HPG axis may alter the gut microbiome directly, through action of sex steroids on the gut microbiome, or indirectly, through sex-specific host physiology that interacts with the gut microbiota. It is notable that the mechanisms that give rise to sex differences in the gut microbiome remain almost completely unknown. To further tease out these mechanisms, future studies should investigate taxonomic and functional sex differences within intestinal microbial communities using metagenomics and metabolomics. Furthermore, the effect of sex steroids and bile acids on growth of individual gut microbes and communities should be investigated to determine if these compounds are sufficient to alter community composition and function in a sex-dependent manner. Additionally, in vivo mouse models of innate or adaptive immune function could be used to determine the contribution of the immune system to sexual maturation of the gut microbiota.

### Implications for microbiome-based therapeutics and future directions

These findings have important implications for treatment of diseases linked to the gut microbiome that have sex bias, alterations in sex steroid levels, and/or develop during specific life stages including childhood (pre-puberty), puberty, and adulthood (post-puberty). As the FDA approved two fecal microbiome products in 2023 for treatment of recurrent *Clostridium difficile*, there is considerable interest in developing novel pro- and postbiotic therapies for many diseases. Given our results, patients may respond to microbiome-based therapeutics differently based on pubertal status, sex steroid levels (e.g., pre-vs. post-menopausal women, women with polycystic ovary syndrome), or medications the patient is taking (e.g. hormone replacement therapy, hormonal contraceptives, gonadotropin-releasing hormone agonist/antagonist).

Additionally, given HPG-dependent sex differences in microbial species diversity, community structure, and metabolites between adult wild-type male and female mice, microbiome-based therapeutics for peri-pubertal and adult patients may need to target different microbial species in men and women.

Our results showing lower connectiveness in adult female wild-type networks also prompts questions concerning gut microbiome community stability in adult women. Given the differences in length between the murine estrous cycle (∼4-5 days) and the human menstrual cycle (∼28 days), studies should investigate whether the menstrual cycle induces similar community instability in women, and whether a woman’s cycle length impacts network stability. This could have implications for microbiome-linked conditions where menstrual cycle length is affected, such as PCOS and endometriosis [15, 81]. It should also be determined whether conditions resulting in static steroid levels, e.g. taking hormonal contraceptives, hormone replacement therapy, or menopause, mirror the hypogonadal state in mice, to determine whether the reduced microbial network connectiveness is related to the oscillating nature of sex steroid levels in the estrous cycle. This could also be tested mechanistically in mice using gonadectomy and steroid replacement studies.

## DATA AVAILABILITY

Sequence reads from this study were deposited into SRA under BioProject PRJNA983444.

## ACKNOWLEDGEMENTS

This work was supported by NIH R01 HD095412 to V.G.T and S.T.K. L.S-H. was funded by NIH F31 HD105403, the Rees Stealy Research Foundation, the San Diego Chapter of the ARCS Foundation, and the SDSU Graduate Division C.O.R.E. Fellowship.

## Notes

### Competing Interest Statement

The authors have declared no competing interest.

