## Supplemental Figures and Tables for "Genetic hypogonadal (Gnrh1^hpg^) mouse model uncovers influence of reproductive axis on maturation of the gut microbiome during puberty"

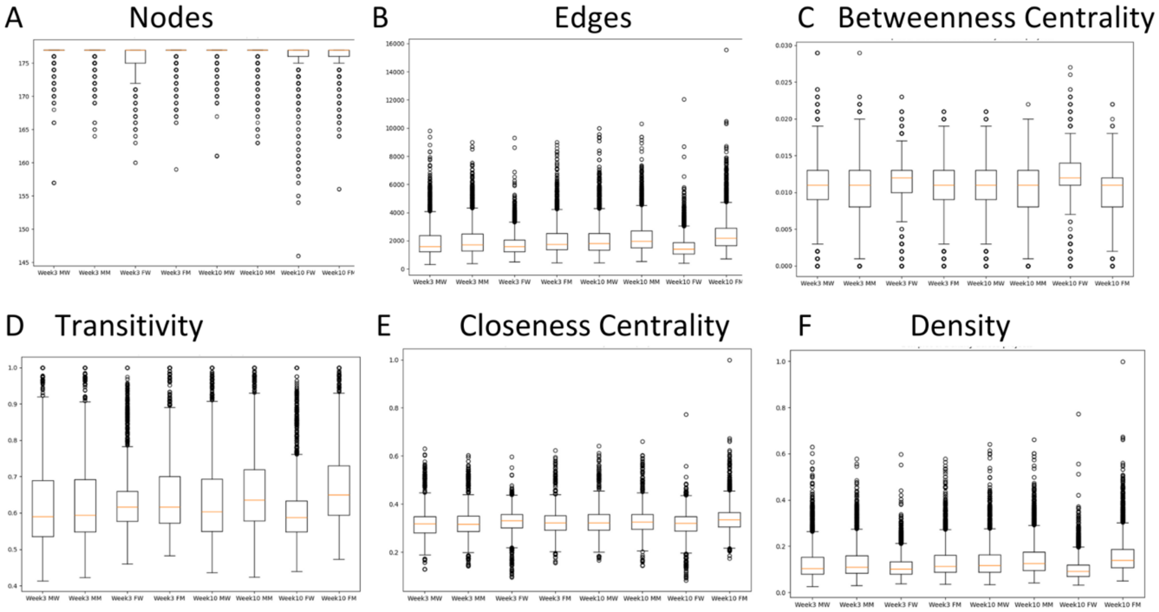


Supplemental Figure 1. Network statistics box and whisker plots for week 3 and week 10 networks.
Networks were computed on 5000 bootstrap replicates of MAG species assignments. (abbreviations: MW = male wild type, MM = male mutant, FW = female wild type, FM = female mutant).

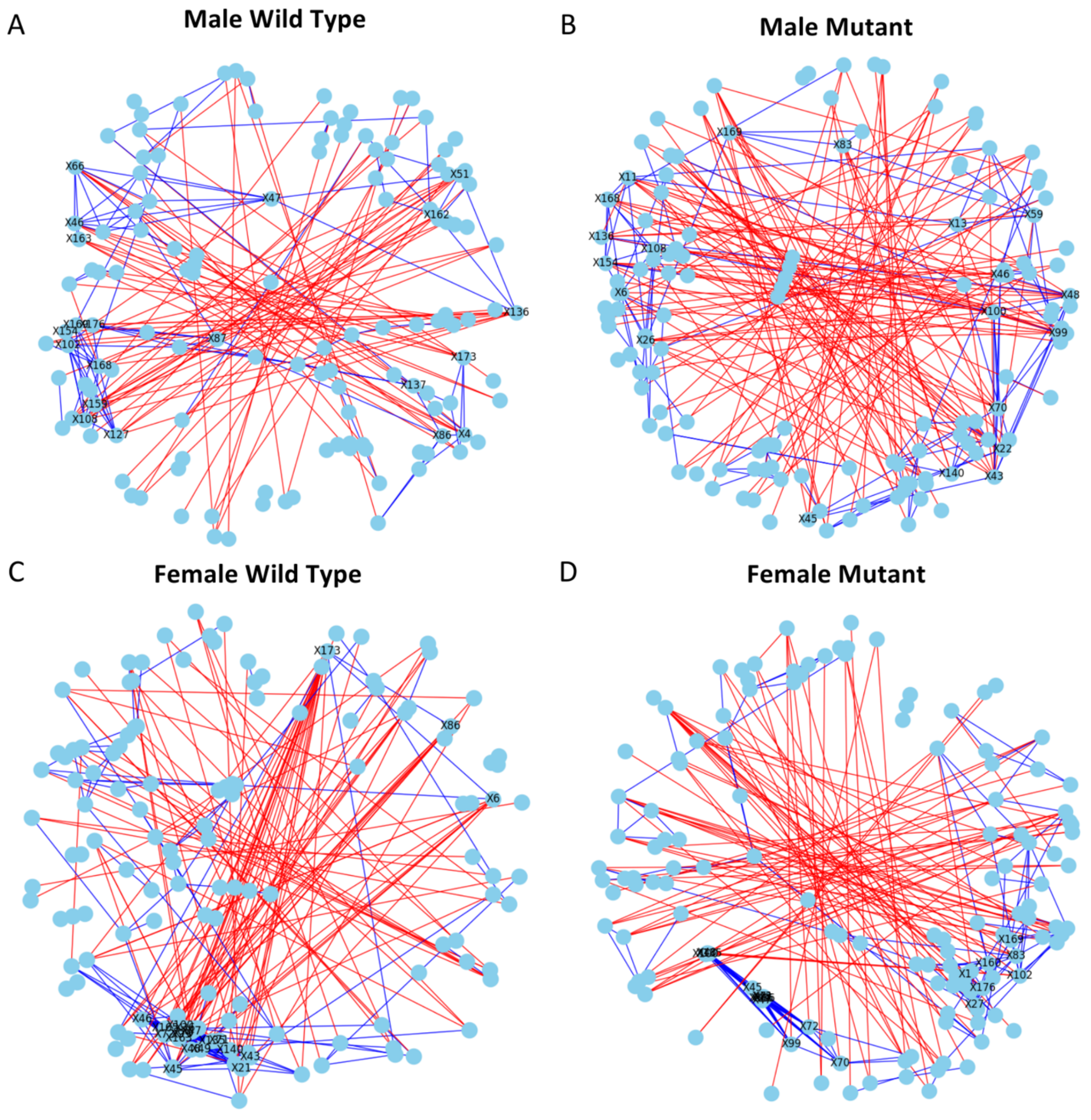


Supplemental Figure 2. Network visualizations for week 3 networks computed on 5000 bootstrap replicates of MAG species assignments.

Networks for (A) male wild-type (B) male mutant (C) female wild-type, and (D) female mutant mice. Blue lines are positive correlation and red lines are negative correlation. Nodes are colored in light blue and represent species. The top 20 species with the most nodes are labeled with numbers corresponding to species listed in Supplemental Table 1.

**Supplemental Tables**

Supplemental Table 1. List of the species indicated by each label in the network plots

| | Species | | Species | | Species |
|---|---|---|---|---|---|
| **X1** | 14-2_sp | **X61** | Coprobacter_secundus | **X120** | MGBC163016_sp910589135 |
| **X2** | 14-2_sp000403315 | **X62** | Coproplasma_sp | **X121** | MGBC163490_sp910588075 |
| **X3** | 14-2_sp004793545 | **X63** | Coproplasma_sp013316055 | **X122** | MGBC164599_sp |
| **X4** | 14-2_sp910576385 | **X64** | Coproplasma_sp910577555 | **X123** | MGBC164599_sp910575625 |
| **X5** | 14-2_sp910577785 | **X65** | Coproplasma_sp910578145 | **X124** | MGBC165282_sp910589265 |
| **X6** | 14-2_sp910580015 | **X66** | Coproplasma_sp910586065 | **X125** | Massilicoli_timonensis |
| **X7** | 1XD42-69_sp | **X67** | D16-34_sp910575165 | **X126** | Merdisoma_sp |
| **X8** | 1XD42-69_sp009911505 | **X68** | D16-63_sp | **X127** | Merdisoma_sp910574475 |
| **X9** | 1XD42-69_sp011959925 | **X69** | Duncaniella_dubosii | **X128** | Mucispirillum_schaedleri |
| **X10** | 1XD8-76_sp910573755 | **X70** | Duncaniella_freteri | **X129** | Muribaculum_gordoncarteri |
| **X11** | Acetatifactor_sp | **X71** | Duncaniella_muris | **X130** | Muribaculum_intestinale |
| **X12** | Acetatifactor_sp011959105 | **X72** | Duncaniella_sp | **X131** | Nanosyncoccus_sp |
| **X13** | Acetatifactor_sp910585425 | **X73** | Dysosmobacter_sp | **X132** | Odoribacter_sp910589025 |
| **X14** | Acetatifactor_sp910585615 | **X74** | Dysosmobacter_sp022797935 | **X133** | Oscillospiraceae_sp |
| **X15** | Acetatifactor_sp910585665 | **X75** | Dysosmobacter_sp910584995 | **X134** | Paralachnospira_sp022792335 |
| **X16** | Acetatifactor_sp910589035 | **X76** | Dysosmobacter_sp910588175 | **X135** | Paramuribaculum_intestinale |
| **X17** | Acutalibacter_sp | **X77** | Enterenecus_sp | **X136** | Paramuribaculum_sp001689565 |
| **X18** | Acutalibacter_sp910580045 | **X78** | Enterenecus_sp910585265 | **X137** | Pelethenecus_sp910577745 |
| **X19** | Acutalibacter_sp910587995 | **X79** | Enterococcus_faecalis | **X138** | Pelethomonas_sp |
| **X20** | Adlercreutzia_mucosicola | **X80** | Enterorhabdus_sp009911695 | **X139** | Pelethomonas_sp910579965 |
| **X21** | Adlercreutzia_muris | **X81** | Eubacterium_F_sp | **X140** | Phocaeicola_barnesiae |
| **X22** | Adlercreutzia_sp | **X82** | Eubacterium_G_sp910574895 | **X141** | Pseudobutyricicoccus_sp910579575 |
| **X23** | Alistipes_sp910575495 | **X83** | Eubacterium_J_sp | **X142** | Pumilibacter_sp |
| **X24** | Alloprevotella_sp002933955 | **X84** | Eubacterium_J_sp910574915 | **X143** | RGIG4057_sp910577675 |
| **X25** | Anaerocaecibacter_sp910577995 | **X85** | Eubacterium_J_sp910579075 | **X144** | RGIG7193_sp |
| **X26** | Anaerotignum_sp910576545 | **X86** | Eubacterium_J_sp910586445 | **X145** | RGIG7193_sp022791755 |
| **X27** | Anaerotruncus_colihominis | **X87** | Eubacterium_plexicaudatum | **X146** | RGIG8607_sp910588535 |
| **X28** | Anaerotruncus_sp | **X88** | Evtepia_sp | **X147** | Roseburia_sp910587865 |
| **X29** | Anaerotruncus_sp000403395 | **X89** | Evtepia_sp910584805 | **X148** | Ruminiclostridium_E_sp910585505 |
| **X30** | Anaerotruncus_sp003612625 | **X90** | Faecivivens_sp022793675 | **X149** | Schaedlerella_sp |
| **X31** | Anaerotruncus_sp910586875 | **X91** | Fimisoma_sp | **X150** | Sporofaciens_sp |
| **X32** | Angelakisella_sp | **X92** | Flavonifractor_plautii | **X151** | Sporofaciens_sp910575835 |
| **X33** | Angelakisella_sp022795575 | **X93** | Gallimonas_sp910586825 | **X152** | UBA11774_sp022790905 |
| **X34** | Angelakisella_sp910585535 | **X94** | HGM11386_sp022791055 | **X153** | UBA1381_sp |
| **X35** | Avidehalobacter_sp022797295 | **X95** | HGM13010_sp022800755 | **X154** | UBA3282_sp |
| **X36** | Bacteroides_acidifaciens | **X96** | JAAVYD01_sp022790955 | **X155** | UBA3282_sp003611805 |
| **X37** | Bacteroides_sp002491635 | **X97** | JAAVYR01_sp022791255 | **X156** | UBA3282_sp009774655 |
| **X38** | Bacteroides_uniformis | **X98** | JAAWBF01_sp | **X157** | UBA3282_sp910575245 |
| **X39** | Butyribacter_sp009774235 | **X99** | JAFLTL01_sp | **X158** | UBA3282_sp910577735 |
| **X40** | Butyricicoccaceae_sp | **X100** | JAFLTL01_sp910578395 | **X159** | UBA3282_sp910578115 |
| **X41** | Butyricimonas_virosa | **X101** | JANFXS01_sp022791155 | **X160** | UBA3402_sp022792815 |
| **X42** | C-19_sp | **X102** | Kineothri_sp | **X161** | UBA7109_sp910575605 |
| **X43** | CAG-485_sp | **X103** | Kineothrix_sp011958945 | **X162** | UBA7173_sp |
| **X44** | CAG-485_sp002361215 | **X104** | Kineothrix_sp910588855 | **X163** | UBA7173_sp001689485 |
| **X45** | CAG-485_sp002362485 | **X105** | Lachnoclostridium_B_sp | **X164** | UMGS1071_sp022801595 |
| **X46** | CAG-873_sp | **X106** | Lachnoclostridium_pacaense | **X165** | Unclassified_BinA |
| **X47** | CAG-873_sp002490635 | **X107** | Lachnoclostridium_phocaeense | **X166** | Unclassified_BinB |
| **X48** | CAG-873_sp009775265 | **X108** | Lachnospiraceae_sp | **X167** | Unclassified_BinC |
| **X49** | CAG-873_sp011959565 | **X109** | Lactobacillus_amylovorus | **X168** | Unclassified_BinD |
| **X50** | CAG-95_sp000403495 | **X110** | Lactobacillus_johnsonii | **X169** | Unclassified_BinE |
| **X51** | CAJFPI01_sp022801715 | **X111** | Lactobacillus_ultunensis | **X170** | Unclassified_BinF |
| **X52** | CAJTFG01_sp910575705 | **X112** | Lawsonibacter_sp | **X171** | Unclassified_BinG |
| **X53** | COE1_sp | **X113** | Lawsonibacter_sp022801435 | **X172** | Unclassified_BinH |
| **X54** | COE1_sp000403215 | **X114** | Ligilactobacillus_murinus | **X173** | Ventrimonas_sp |
| **X55** | COE1_sp003513705 | **X115** | MD308_sp910584615 | **X174** | Ventrimonas_sp910577765 |
| **X56** | COE1_sp910585765 | **X116** | MGBC131033_sp910585655 | **X175** | Zhenpiania_sp |
| **X57** | Caccovicinus_sp910575565 | **X117** | MGBC136627_sp910585935 | **X176** | [Clostridium]_clostridioforme |
| **X58** | Choladocola_sp | **X118** | MGBC163016_sp | **X177** | [Clostridium]_innocuum |
| **X59** | Choladocola_sp009774145 | **X119** | MGBC163016_sp910588905 | | |
| **X60** | Clostridium_Q_sp | **X120** | MGBC163016_sp910589135 | | |

Supplemental Table 2. Taxonomic identities of the 20 most-connected nodes (species) for each week 3 network. E = Edges

| **Male Wild-Type Week 3** | | **Male Mutant Week 3** | | **Female Wild Type Week 3** | | **Female Mutant Week 3** | |
|---|---|---|---|---|---|---|---|
| **Species** | | **Species** | | **Species** | **E** | **Species** | **E** |
| Coproplasma_ sp910586065 (Borkfalkiaceae) | 11 | CAG-873_sp009775265 (Provetellaceae ) | 14 | CAG-873_ sp009775265 (Provetellaceae ) | 21 | Duncaniella_sp (Muribaculaceae) | 15 |
| Paramuribaculum sp001689565 (Muribaculaceae) | 11 | UBA3282_sp (Lachnospiraceae) | 13 | CAG-873_ sp011959565 (Provetellaceae ) | 21 | JAFLTL01_sp (Muribaculaceae) | 13 |
| Unclassified_BinD | 11 | CAG-485_sp (Provetellaceae) | 13 | CAG-873_sp (Muribaculaceae ) | 20 | [Clostridium] clostridioforme (Lachnospiraceae) | 12 |
| CAG-873_sp (Provetellaceae ) | 10 | Acetatifactor_sp (Lachnospiraceae) | 12 | Ventrimonas_sp (Lachnospiraceae) | 19 | Anaerotruncus_ colihominis (Oscillospiraceae) | 12 |
| Kineothrix_sp (Lachnospiraceae) | 10 | JAFLTL01_sp910578395 (Muribaculaceae) | 12 | Adlercreutzia_muris (Eggerthellaceae) | 19 | Duncaniella_freteri (Muribaculaceae) | 12 |
| Unclassified_BinE | 10 | Adlercreutzia_sp (Eggerthellaceae) | 11 | CAG-873_sp002490635 ( Provetellaceae ) | 19 | CAG-485_sp ( Provetellaceae ) | 12 |
| Lachnospiraceae_sp | 10 | Duncaniella_freteri (Muribaculaceae) | 11 | JAFLTL01_sp (Muribaculaceae) | 19 | JAFLTL01_sp (Muribaculaceae) | 12 |
| Merdisoma_ sp910574475 (Lachnospiraceae) | 10 | JAFLTL01_sp (Muribaculaceae) | 11 | UBA7173 _sp001689485 (Muribaculaceae) | 19 | Paramuribaculum_ intestinale | 12 |
| [Clostridium]_ clostridioforme (Lachnospiraceae) | 9 | Anaerotignum_ sp910576545 (Anaerotignaceae) | 10 | JAFLTL01_sp (Muribaculaceae) | 19 | Paramuribaculum_ sp001689565 | 12 |
| CAJFPI01_ sp022801715 (Oscillospiraceae) | 9 | Unclassified_BinD | 9 | Duncaniella_sp (Muribaculaceae) | 18 | CAG-873_sp (Provetellaceae ) | 12 |
| Pelethenecus_ sp910577745 | 8 | CAG-873_sp ( Provetellaceae ) | 9 | Unclassified_BinA | 17 | CAG-873_sp002490635 ( Provetellaceae ) | 12 |
| CAG-873_ sp002490635 ( Provetellaceae ) | 8 | Choladocola_sp009774145 (Bartonellaceae) | 8 | Phocaeicola_barnesiae (Bacteroidaceae) | 16 | CAG-873_sp009775265 ( Provetellaceae ) | 12 |
| UBA3282_ sp910578115 (Unidentified Lachnospiraceae) | 8 | 14-2_sp910580015 ( Lachnospiraceae) | 8 | Duncaniella_muris (Muribaculaceae) | 16 | CAG-873_sp011959565 ( Provetellaceae ) | 12 |
| 14-2_ sp910576385 ( Lachnospiraceae) | 7 | Lachnospiraceae_sp | 8 | Paramuribaculum Intestinale (Muribaculaceae) | 14 | Duncaniella_muris ( Muribaculaceae) | 12 |
| Ventrimonas_sp (Lachnospiraceae) | 7 | Unclassified_BinE | 8 | Duncaniella_freteri (Muribaculaceae) | 14 | Eubacterium_J_sp ( Eubacteriaceae) | 11 |
| UBA7173_sp ( Muribaculaceae) | 7 | Acetatifactor_ sp910585425 | 7 | CAG-485_sp ( Provetellaceae ) | 13 | Unclassified_BinE | 11 |
| UBA3282_sp (Unidentified Lachnospiraceae) | 7 | Paramuribaculum_ sp001689565 (Muribaculaceae) | 7 | CAG-485_sp002362485 ( Provetellaceae ) | 12 | CAG-485_sp002362485 ( Provetellaceae ) | 11 |
| UBA7173_ sp001689485 ( Muribaculaceae) | 6 | Phocaeicola_barnesiae (Bacteroidaceae) | 7 | 14-2_sp910580015 ( Lachnospiraceae) | 10 | Kineothrix_sp (Lachnospiraceae) | 10 |
| Eubacterium_J_ sp910586445 (Lachnospiraceae) | 6 | CAG-485_sp002362485 ( Provetellaceae ) | 7 | Bacteroides_acidifaciens (Bacteroidaceae) | 9 | 14-2_sp (Lachnospiraceae) | 9 |
| Eubacterium_ plexicaudatum (Eubacteriaceae) | 6 | Eubacterium_J_sp ( Lachnospiraceae) | 7 | Eubacterium_Jsp910586445 ( Lachnospiraceae)) | 9 | UBA3402_sp022792815 (Unidentified Lachnospiraceae) | 9 |

Supplemental Table 3. Taxonomic identities of the 20 most-connected nodes (species) for each week 10 network. E = Edges

| Male Wild-Type Week 10 | | Male Mutant Week 10 | | Female Wild Type Week 10 | | Female Mutant Week 10 | |
|---|---|---|---|---|---|---|---|
| **Species** | **E** | **Species** | **E** | **Species** | **E** | **Species** | **E** |
| UBA7173_sp001689485 ( Muribaculaceae) | 17 | CAG-873_sp (Provetellaceae ) | 17 | JAFLTL01_sp (Muribaculaceae) | 18 | Kineothrix_sp (Lachnospiraceae) | 24 |
| CAG-873_sp (Provetellaceae ) | 14 | Duncaniella_muris (Unidentified Muribaculaceae) | 17 | Enterorhabdus_ sp009911695 (Eggerthellaceae) | 16 | UBA7173_sp (Muribaculaceae) | 23 |
| Duncaniella_freteri (Muribaculaceae) | 14 | Duncaniella_sp (Unidentified Muribaculaceae) | 17 | Adlercreutzia_sp (Eggerthellaceae) | 15 | CAG-873_sp002490635 (Provetellaceae ) | 22 |
| CAG-485_sp002361215 (Provetellaceae ) | 13 | Muribaculum_intestinale (Muribaculaceae) | 16 | CAG-873_sp002490635 (Provetellaceae ) | 15 | Evtepia_sp (Eubacteriales) | 22 |
| CAG-873_sp002490635 ( Provetellaceae ) | 12 | CAG-485_sp002361215 ( Provetellaceae ) | 16 | CAG-873_sp011959565 ( Provetellaceae ) | 14 | Dysosmobacter (Oscillospiraceae) | 22 |
| Duncaniella_sp (Muribaculaceae) | 12 | Caccovicinus_ sp910575565 (Lachnospiraceae) | 14 | CAG-873_sp (Provetellaceae ) | 14 | Coprobacter_secundus (Barnesiellaceae) | 20 |
| Duncaniella_muris (Muribaculaceae) | 11 | CAG-485_sp002362485 (Provetellaceae ) | 14 | CAG-873_sp009775265 (Provetellaceae ) | 14 | JAAVYD01_sp022790955 (Anaerovoracaceae) | 17 |
| Lachnospiraceae_sp | 11 | JAAVYD01_sp022790955 (Anaerovoracaceae) | 13 | Duncaniella_muris (Unidentified Muribaculaceae) | 14 | Unclassified_BinD | 17 |
| Acetatifactor_sp910585425 (Lachnospiraceae) | 10 | Lachnospiraceae_sp | 13 | JAFLTL01_sp (Muribaculaceae) | 14 | Angelakisella_sp022795575 (Ruminococcaceae) | 17 |
| Evtepia_sp910584805 (Eubacteriales) | 10 | Unclassified_BinG | 13 | Lawsonibacter_sp (Oscillospiraceae) | 14 | Lactobacillus amylovorus (Lactobacillaceae) | 17 |
| UBA7173_sp ( Muribaculaceae) | 10 | Schaedlerella_sp (Lachnospiraceae) | 13 | Duncaniella_sp (Unidentified Muribaculaceae) | 12 | Merdisoma_sp (Lachnospiraceae) | 17 |
| 1XD8-76_sp910573755 ( Lachnospiraceae) | 10 | Coprobacter_secundus (Barnesiellaceae) | 12 | UBA7173_sp001689485 (Muribaculaceae) | 12 | RGIG7193_sp (Lachnospiraceae) | 17 |
| JAFLTL01_sp (Muribaculaceae) | 10 | CAG-485_sp (Provetellaceae ) | 12 | D16-34_sp910575165 (Eggerthellaceae) | 11 | Roseburia_sp910587865 (Lachnospiraceae) | 17 |
| CAG-873_sp009775265 (Provetellaceae ) | 10 | Eubacterium_G_ sp910574895 | 12 | Duncaniella_freteri (Unidentified Muribaculaceae) | 11 | MGBC165282_ sp910589265 (Lachnospiraceae) | 16 |
| Evtepia_sp ( Eubacteriales) | 10 | 14-2_sp004793545 (Lachnospiraceae) | 11 | Acetatifactor_sp (Lachnospiraceae) | 10 | CAG-485_sp002361215 (Provetellaceae ) | 16 |
| Lachnoclostridium_pacaense (Lachnospiraceae) | 9 | Duncaniella_freteri (Muribaculaceae) | 11 | Mucispirillum schaedleri (Mucispirillaceae) | 10 | Phocaeicola_barnesiae (Bacteroidaceae) | 15 |
| [Clostridium]_innocuum (Coprobacillaceae) | 9 | Ventrimonas_sp910577765 (Lachnospiraceae) | 10 | 1XD8-76_sp910573755 (Lachnospiraceae) | 7 | JAFLTL01_sp (Muribaculaceae) | 15 |
| Lachnoclostridium_phocaeense (Lachnospiraceae) | 9 | Evtepia_sp910584805 (Eubacteriales) | 10 | Lactobacillus_johnsonii (Lactobacillaceae) | 7 | Duncaniella_freteri (Muribaculaceae) | 15 |
| 14-2_sp004793545 (Lachnospiraceae) | 8 | Lachnoclostridium_B_sp (Lachnospiraceae) | 10 | 14-2_sp910576385 (Lachnospiraceae) | 6 | Clostridium_Q_sp (Lachnospiraceae) | 14 |
| Coprobacter_secundus (Barnesiellaceae) | 8 | Pelethomonas_sp (Oscillospiraceae) | 10 | [Clostridium]_innocuum (Coprobacillaceae) | 6 | Pelethomonas_sp (Oscillospiraceae) | 14 |
